# Met/HGFR triggers detrimental reactive microglia in TBI

**DOI:** 10.1101/2021.12.05.471232

**Authors:** Rida Rehman, Michael Miller, Sruthi Sankari Krishnamurthy, Jacob Kjell, Lobna Elsayed, Florian olde Heuvel, Alison Conquest, Akila Chandrasekar, Albert Ludolph, Tobias Boeckers, Medhanie A Mulaw, Magdalena Goetz, Maria Cristina Morganti-Kossmann, Aya Takeoka, Francesco Roselli

**Affiliations:** Dept. of Neurology, Ulm University, Ulm DE; Neuro-Electronics Research Flanders (NERF), Flanders Institute for Biotechnology (VIB), Leuven, Belgium; Department of Neuroscience and Leuven Brain Institute, KU Leuven, Leuven, Belgium; Institute for Stem Cell Research, Helmholtz Zentrum München & Biomedical Center, Ludwig-Maximilians Universitaet, Planegg-Martinsried, Germany; Department of Clinical Neuroscience, Karolinska Institutet, Solna, Sweden; National Trauma Research Institute and Department of Neurosurgery, The Alfred Hospital, Melbourne, Australia; Dept. of Cellular Neurobiology, Technical University, Braunschweig, Germany; German center for Neurodegenerative Diseases (DZNE)-Ulm; Dept. of Internal Medicine I and Institute of Molecular Medicine, Ulm University, Ulm DE; Department of Epidemiology and Preventive Medicine, Monash University, Melbourne, Australia; IMEC, Leuven, Belgium

**Keywords:** Traumatic Brain Injury, antibody array, proteomics, cMet, HGFR, Btk, VEGFR, microglia, neuroinflammation

## Abstract

The complexity of the signaling events, cellular responses unfolding in neuronal, glial and immune cells upon Traumatic brain injury (TBI) constitutes an obstacle in elucidating pathophysiological links and targets for intervention. We used array phosphoproteomics in a murine mild blunt TBI to reconstruct the temporal dynamics of tyrosine-kinase signaling in TBI and then to scrutinize the large-scale effects of the perturbation of cMet/HGFR, VEGFR1 and Btk signaling by small molecules. cMet/HGFR emerged as a selective modifier of the early microglial response, and cMet/HGFR blockade prevented the induction of microglial inflammatory mediators, of reactive microglia morphology and of TBI-associated responses in neurons, vessels and brain extracellular matrix. Acute or prolonged cMet/HGFR inhibition ameliorated neuronal survival and motor recovery. Early elevation of HGF itself in the CSF of TBI patients suggest that this mechanism has translational value in human subjects. Our findings identify cMet/HGFR as a modulator of early neuroinflammation in TBI with translational potential and indicate several RTK families as possible additional targets for TBI treatment.

**Summary:** Controlling neuroinflammation in neurotrauma is an important but unachieved goal. This study exploits a moderate TBI model and array-based proteomics to identify cMet as a new inducer of reactive microglia. A small-molecule inhibitor of cMet contains microglial reactivity, reduces neuronal and vascular alterations, limits behavioural disturbances and accelerates recovery.

**Highlights:**

- Met is activated in microglia upon TBI and drives microglial reactivity.
- A Met inhibitor reduces motor dysfunction upon TBI and promotes recovery.
- Blockade of MET prevents the appearance of a reactive microglia.
- The cMET inhibitor reduces the sub-acute neuronal loss after TBI.

## Introduction

Traumatic Brain Injury (TBI) is characterized by a highly complex and dynamic set of responses taking place in neurons, glial cells (in particular astrocytes, NG2 glia and microglia), endothelial cells and infiltrating leukocytes (Alam et al., 2020) and unfolding over time, from hours-days (acute phase) to weeks and months (chronic phase). Microglia reactions are among the very first events following TBI; upon injury, microglia cells increase their motility, retract processes, assume a chemotactic morphology and rapidly reach the site of damage (Davalos et al., 2005; Jassam et al., 2017; Nimmerjahn et al., 2005; Roth et al., 2014). A number of detrimental consequences of acute microglial responses have been described, including furthering local acidosis (Ritzel et al., 2021), production of reactive oxygen and nitric oxide radicals (Ma et al., 2018), inducing widespread synaptic silencing (Zhang et al., 2014) and contributing to neurotoxicity (Wang et al., 2020). Taken together, these processes may determine the “secondary damage”, the progressive loss of neurons that follows (for hours to days) the acute lesion caused by the mechanical forces. Nevertheless, the contribution of microglia is considered more nuanced (Jassam et al., 2017) and beneficial effects involve the effective sealing of the disrupted glia limitans (Roth et al., 2014), the rapid clearing of neuronal debris (Herzog et al., 2019) and the stimulation of brain repair through neurogenesis (Willis et al., 2020). The acute-phase neuroinflammatory response to TBI is further complicated by the breakdown of Blood-Brain-Barrier (BBB; O’Keeffe et al., 2020) and by the infiltration of neutrophils, macrophages and lymphocytes (Szmydynger-Chodobska et al., 2012), allowing the extravasation of blood-derived proteases and complement systems (Hopp et al., 2017; Roselli et al., 2018) which contributes to brain oedema (Zhou et al., 2020). Damage Associated Molecular Patterns (DAMPs) are the early drivers of microglia reactivity upon brain damage and include ATP released by damaged cells (Roth et al., 2014; Wicher et al., 2017), interferons, complement factors (Chandrasekar et al., 2018; Roselli et al., 2018) and inflammatory cytokines (Wicher et al., 2017). The large-scale architecture of TBI pathophysiology is therefore critically affected by the quantity and pattern of DAMPS, which in turn depend on the severity of the tissue disruption, the extent of necrosis, haemorrhage, hypoxia and bone lesions. While the majority of cases are scored mild or moderate and display limited tissue damage, even mild TBI can lead to substantial long-term cognitive and behavioural consequences (Wang et al., 2021; Bai et al., 2020).The DAMPS driving microglial reactivity and its consequences at neuronal level in this common TBI setting are only partially understood and the fine-tuning of acute microglial reaction as well as of the overall neuroinflammatory cascade in TBI requires a more comprehensive understanding of the mediators (many of which may still be unknown), regulatory mechanisms and ultimately their impact on long-term consequences.

A substantial number of Receptor and Non-Receptor Tyrosine Kinases (RTK and NRTK, respectively) are abundantly expressed in microglia and endothelial cells with some degree of selectivity (Tondo et al., 2019; Zeisel et al., 2015). Only a few RTK have been shown to have important physiological roles for microglial reactivity in TBI-related conditions: the RTK Axl decreases microglial reactivity in inflammatory conditions and to drive debris phagocytosis (Fourgeaud et al., 2016; Ray et al., 2017) whereas the MCSFR1 (c-Fms, CSFR1) determines microglial survival and proliferation (Elmore et al., 2014; Hu et al., 2020). Nevertheless, the number of RTK expressed in microglia (including interleukin, neurotrophins and growth factors receptors; Wu et al., 2020; Tondo et al., 2019; Zeisel et al., 2015) raises the hypothesis that they may act as DAMP sensors and contribute to early reactivity upon tissue damage.

Importantly, RTK/NRTK display substantial potential as entry points for therapeutic modulation of TBI-induced neuroinflammation, with more than 30 RTK/NRTK inhibitors already in clinical use (Huang et al., 2020) lending themselves to drug repurposing efforts, but it is still largely untapped.

Here we have employed large-scale phosphoproteomic arrays to understand the temporal activation of RTKs and NRTKs in trauma and identify new regulators of TBI-induced neuroinflammation. We have identified the RTK cMet/HGFR as a new player involved in acute microglial activation in TBI. We demonstrate that a small molecule inhibitor of cMet/HGFR is able to prevent inflammatory signaling, limit microglial recruitment, reduce neuronal and vascular distress and acutely improve neurological function upon TBI.

## Results

### Large-scale dynamics of RTK phosphorylation in the cerebral cortex upon TBI

We explored the injury-related architecture of RTK signaling in a murine model of mild TBI. In our weight-drop mild (NSS 0-1; ranging between 0 and 2; Chandrasekar et al., 2018) blunt head trauma model, neuronal counts were still normal 3h after trauma (Figure S1B, C) but significantly declined in the injury core at 3dpi (21.1±15.4% of NeuN+ cells lost in the core lesion-layer II-III; Figure S1A, p=0.0103 vs sham; Figure S1B, C) and neurons were strongly depopulated at 7 dpi (95.5±3.3% in the core; p<0.0001 vs sham). Reciprocally, GFAP+ astrocytes increased at 3 (261.9%±64.2; p= 0.0052 vs sham) and 7 dpi (766.4± 406%; p<0.0001 vs sham; Figure S1B and 1D).

In this model we first monitored, using chemiluminescence-based nitrocellulose antibody arrays, the phosphorylation of 39 distinct RTK in cortical samples obtained 3h (n=7), 1d (n=6), 3d (n=6) or 7d (n=6) post-injury compared with sham controls (n=6) (Figure 1A). After quality control analysis (indicated in the methods), Principal component analysis (PCA) revealed a clear separation of the sham, 3h, 1 and 3 dpi samples, whereas 7 dpi samples displayed substantial variability (Figure 1B). A distinct pattern of RTK phosphorylation for each timepoint considered was detected using modified differential expression analysis algorithms (Figure 1C, D, see also Table S1).

**Figure 1.**
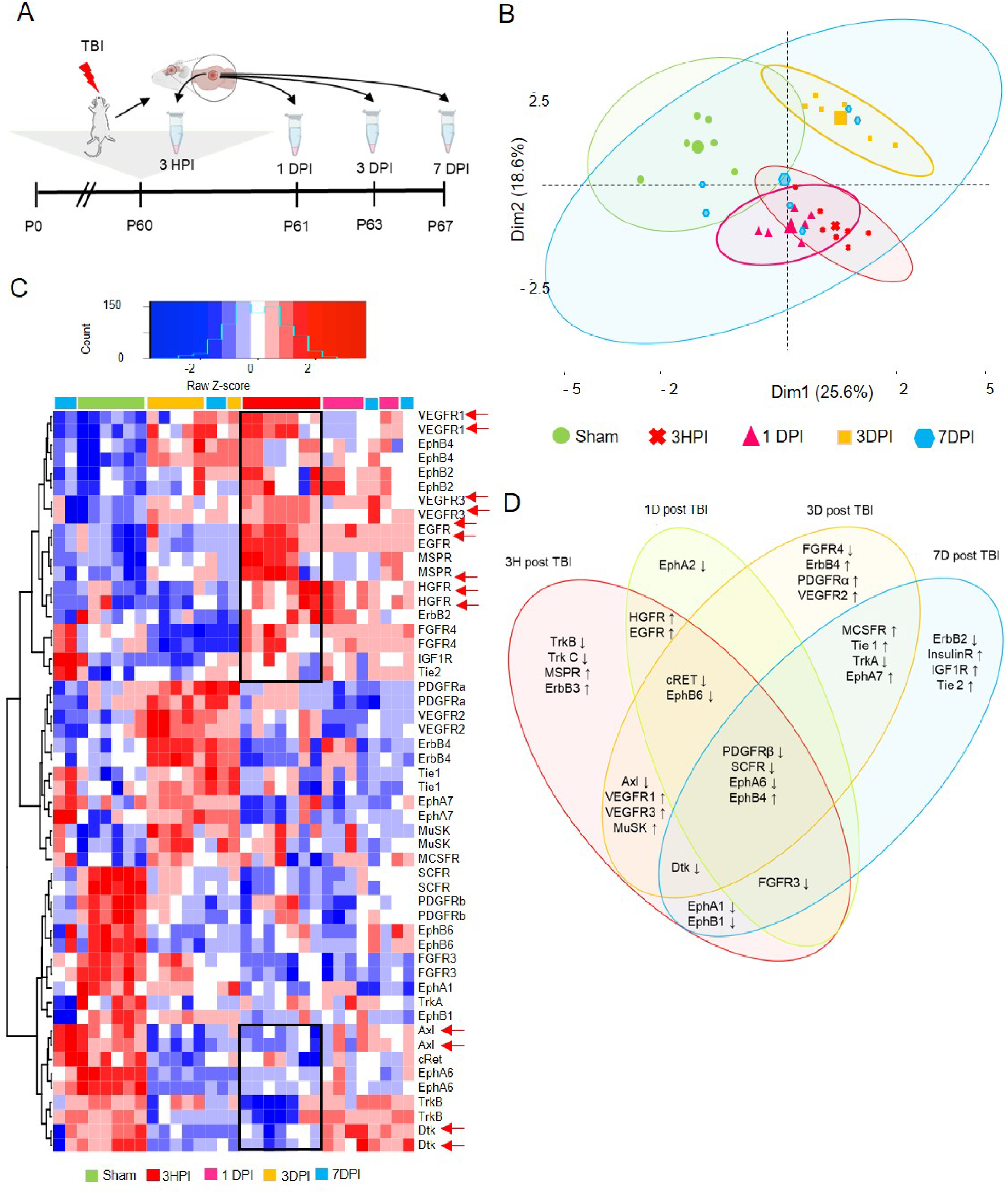
Dynamic RTK phosphorylation pattern post trauma. (A) Schematic outline of the experiment shows mice were subjected to trauma and samples were collected at 3h, 1d, 3d and 7d post injury (DPI) timepoints to map the phosphorylation pattern. (B) Post preprocessing principal component analysis (PCA) plot showing distinct clustering of each group with 95% confidence ellipses around the groups. All groups show minimal overlap compared to the 7DPI group displaying substantial variability. (C) Heatmap of proteins showing unsupervised clustering of differentially expressed (DE) RTKs at different timepoints. The arrows highlight prominent receptors VEGFR1, VEGFR3, HGFR and EGFR. (D) Venn diagram summarizing the overlap of significant RTKs at different timepoints indicating increase or decrease in phosphorylation levels compared to sham. In (B)-(D), n=6 for Sham, 1DPI, 3DPI and 7dpi, and n=7 for 3HPI. Significance for DE proteins was set at P<0.05 (FDR adjusted). Detailed individual comparisons between respective groups and the overlapping phosphorylation pattern for the proteins are shown in Table S1. Also see Figure S1.

The RTK phosphorylation profiles at 3h and 1dpi displayed the largest divergence from the baseline (sham), but were otherwise similar to each other. Statistically significant increase in phosphorylation levels of VEGFR1, VEGFR3, EphB4, cMet/HGFR, MSPR, EGFR, ErbB3, and MuSK was detected at 3h post injury (3HPI). On the other hand, significant down-phosphorylation was seen for TrkB, TrkC, Axl, Dtk, SCFR/c-kit, PDGFRb, cRET, EphA1, EphA6, EphB1 and EphB6. A notable difference in RTK phosphorylation profile was observed at 3dpi, characterized by the significant increase in phosphorylation levels of ErbB4, PDGFRa and VEGFR2 and down phosphorylation of FGFR4, Axl, cRet. Despite the substantial variability observed at 7 dpi, possibly indicating an inter-individual divergence in the sub-acute phase, we detected a statistically significant increase in phosphorylation levels of insulin receptor1, Tie2 and IGFR1 and down-phosphorylation in ErbB2 (Figure 1D).

Among the RTK activated at 3h, at least six are involved in regulation of microglial reactivity: Axl, Dtk and c-kit/SCFR (when activated, they restrain microglial reactivity Fourgeaud et al., 2016; Terashima et al., 2018; Figure S1E) are de-activated whereas the Hepatocyte Growth Factor Receptor Family members (cMet/HGFR and MSPR, involved in chemotaxis (Moransard et al., 2010; Di Renzo et al., 1993), are strongly (but transiently) activated (Figure S1F).

Other RTK families follow unique dynamics upon TBI: phospho-VEGFR1 and phospho-VEGFR3 are significantly upregulated at 3h (strong trend for VEGFR3) and 3 dpi whereas VEGFR2 was only significantly upregulated at 3 dpi (Figure S1F). Significant and time-dependent increase in phosphorylation was observed also for receptors in the EGFR/ErbB family, Insulin/IGF IR family and most notably in the Ephrin family members. EGFR phosphorylation levels were increased at 3h and 1dpi, but returned then to baseline, whereas ErbB3 phosphorylation levels were increased at 3h and 3dpi and ErbB4 was activated only at 3dpi (Figure 1D, Figure S1H). Insulin receptor family (Insulin R and IGFR1) showed a significant increase in phosphorylation levels only at 7 dpi (Figure S1H).

Notably, the RTK phosphorylation pattern was influenced by the severity of the injury: samples from a milder trauma (total energy = 0.47J), when compared to those from standard trauma (total energy = 0.53J) displayed a phosphorylation pattern intermediate between sham and standard-trauma samples (Figure S1I). Interestingly, the milder trauma samples displayed an increased, rather than decreased, phosphorylation of Dtk and Axl, whereas phosphorylation levels of cMet/HGFR and VEGFR3 were still significantly upregulated (Figure S1I). Thus, upon mild TBI, RTK phosphorylation profile is highly dynamic, displaying substantial changes over time and depending on the severity of the trauma.

### In-depth characterization of tyrosine-kinase signaling network upon acute TBI

In order to confirm and extend the data from the chemiluminescence arrays, and to identify signaling cascades downstream of activated RTK, we profiled the tyrosine phosphorylation of 228 distinct target in somatosensory cortex of sham-operated (n=3) or TBI (n=3) mice at the 3h time point after trauma using a fluorescence antibody array. After quality controls (indicated in the methods), Principal Component Analysis (PCA) demonstrated a remarkable separation of sham and TBI ellipses (95% confidence interval; Figure 2A). As depicted in the Volcano plot (Figure 2B), when a cutoff value of 2*log10^2^ was applied, 39 proteins showed statistically significant differences in tyrosine phosphorylation upon trauma (Figure 2C, see also Table S2), including a number of RTK (VEGFR1, EGFR and cMet/HGFR) already revealed by the chemiluminescence screening (cfr. Figure 1C,D and Figure 2C). Notably, several NRTK and other signaling molecules whose phosphorylation was altered upon TBI (such as Btk, Hck, DAPP1, Blnk, Vav1 and Plcg2) are enriched and highly involved in microglial function (Nam et al., 2018; Shah et al., 2009; Sierksma et al., 2020; Suh et al., 2005), whereas a smaller set of hits was attributed to neuronal (NMDAR1 NR2A/2B, Kv1.3 and alpha-Synuclein) origin while for a number of hits, such as cortactin, actin and caspase 9, expression is broad (Figure 2C). We confirmed the cellular specificity of the phosphorylation of three selected targets (prioritized according to the availability of selective inhibitors with good brain penetration and potential for drug repurposing), namely phospho cMet/HGFR (Tyr1234), phosphoBTK (Tyr222); with an anticipated profile of microglial expression, and phosphoVEGFR1 (Tyr1333); with a broader predicted expression profile. Immunoreactivity for phosphorylated cMet/HGFR and pBTK was almost undetectable in sham brains, but was significantly increased upon TBI; notably, nearly all immunoreactivity was restricted to Iba1+ cells (Figure 2D, E, p-cMet/HGFR; p<0.0001 vs sham and pBtk; p=0.0001 vs sham). Neurons displayed very weak immunostaining for phospho-cMet/HGFR and phospho-Btk with no increase in phosphorylation after trauma (Figure S2A, B). On the contrary, immunoreactivity for phospho-VEGFR1 was upregulated in vessels, microglia and neurons (Figure 2D,E, pVEGFR1; microglia: p=0.0044 vs sham, neurons: p=0.0128 vs sham, Figure S2C,D).

**Figure 2.**
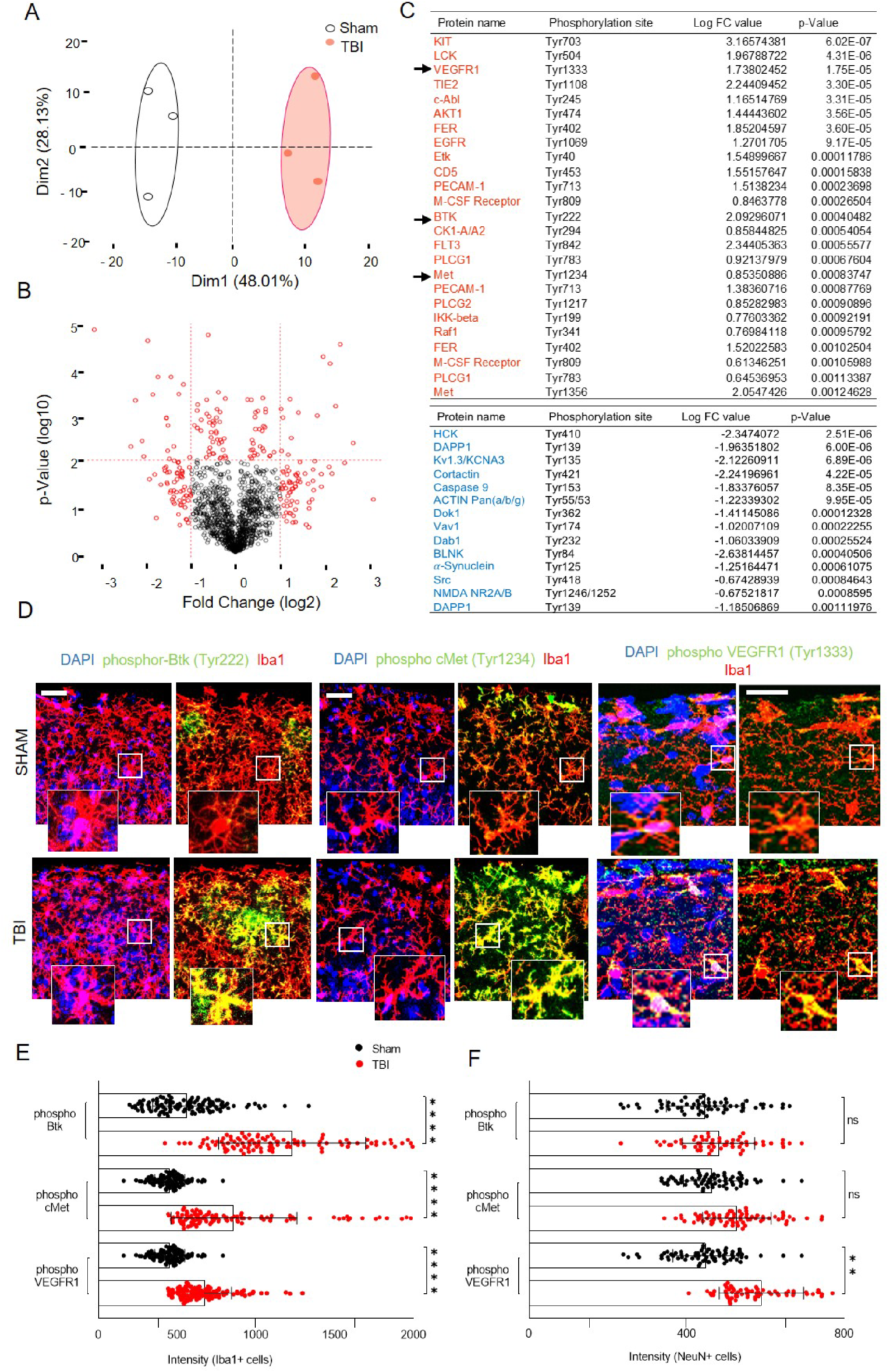
In-depth tyrosine kinase screening 3h post injury and identifying microglia as key player. (A) Post preprocessing principal component analysis (PCA) plot showing distinct clustering of groups with no overlap between the confidence ellipses (95%) with PC1 contributing to 48%. (B) Volcano plot showing data from tyrosine kinase screening of the injured cortex compared to sham at 3h post injury. Cutoff was applied at 2*log10^2^. (C) List of DE proteins (P ≤ 0.05) up-phosphorylated (red) or down-phosphorylated (blue) at 3 hours post trauma compared to baseline (TBI vs Sham) with logarithmic fold change and individual significance. (D) Immunostaining of phospho-Btk, phospho-cMet/HGFR and phospho-VEGFR1 showing colocalization with microglia (Iba1+ cells) and upregulation in TBI compared to sham. (See also Figure S2 for neuronal and vascular overlap of the phosphorylated proteins). (E)-(F) The bar graph showing the intensity levels of phosphorylated receptors expressed in (E) Iba1+ cells (microglia) or (F) NeuN+ cells (neurons) verifying the screening results. All graphs are represented as mean ± SD. In (A)-(C), n=3 for Sham and TBI. Significance for DE proteins was set at P<0.05 (BH adjusted). Detailed individual comparisons between TBI and Sham groups are shown in Table S2. In (D)-(F), n= 4 for each staining. Dots indicate individual values for each cell. Significance of differences between means were analyzed using two-way anova test with Sidak correction (recommended). (ns= not significant, **P=0.0044, ***P<0.0001). Scale bars: 200µm for (D).

### Small-molecule inhibitors of cMet/HGFR and VEGFR substantially alter the signaling architecture upon TBI

We then focused on exploring the functional relevance of Btk, cMet/HGFR and VEGFR signaling on TBI-induced microglial activation, using small-molecule kinase inhibitors. Although small-molecules targeting kinases are less selective than gene-knockout manipulation, they lend themselves to explore acute effects (i.e., do not suffer from life-long target loss and unforeseen adaptation) and display stronger translational potential. Based on the selectivity and pharmacokinetic data available (in particular, blood-brain-barrier penetration), we selected the Btk inhibitor Spebrutinib (CC-292; IC50 of <0.5 nM, displaying at least 1400-fold selectivity over the other kinases; Evans et al., 2013; Tonge, 2018), the cMet/HGFR inhibitor JNJ-38877605 (inhibitor of c-Met with IC50 of 4 nM, 600-fold selective for c-Met than 200 other tyrosine and serine-threonine kinases; De Bacco et al., 2016; Etnyre et al., 2014; Lolkema et al., 2015) and the broader selectivity VEGFR inhibitor Valatanib (PTK-787; inhibit VEGFR1 and VEGFR2/KDR with IC50 of 77nM and 37 nM, respectively; also inhibits Flk, c-Kit and PDGFRβ with IC50 of 270 nM, 730 nM and 580 nM, respectively; Wood et al., 2000; Kong et al., 2017; Reardon et al., 2009; Shankar et al., 2016).

First, we validated the target engagement of the three inhibitors by administering each molecule (or vehicle alone) (BTK inhibitor: 30mg/kg [Evans et al., 2013], cMet/HGFR inhibitor: 40mg/kg [De Bacco et al., 2016; Lolkema et al., 2015], VEGFR inhibitor: 50mg/kg [Kong et al., 2017; Reardon et al., 2009; Shankar et al., 2016]) 2h before trauma (based on reported pharmacokinetic data) and verifying a significant decrease in p-Btk (Tyr222), p-cMet/HGFR (Tyr1234) and p-VEGFR1 (Tyr1333) 3h after trauma by whole-tissue western blot (Figure 2A-C, see also Figure S3; TBI vs sham: phospho-Btk p=0.0029, phospho-cMet/HGFR p=0.028, phospho-VEGFR p=0.011, TBI vs inhibitor TBI: phospho-Btk p=0.0042, phospho-cMet/HGFR p=0.0065, phospho-VEGFR p=0.015).

Next, we explored the large-scale impact (by phospho-tyrosine antibody arrays) of selective inhibition with single dose (SD) of Btk [Inhibitor TBI (SD Btk), n=4], cMet/HGFR [Inhibitor TBI (SD cMet), n=4] or VEGFR [Inhibitor TBI (SD VEGFR), n=3) upon TBI (3h time point), contrasted against saline-treated TBI (n=4) or sham-operated (n=4) mice to explore the signaling landscape set in motion by TBI. PCA plots of phosphotyrosine array datasets revealed a distinct separation of groups in PCA plots, with dimension 1 contributing to 31.41% of the variability indicating substantial and significant effect of the inhibitors (Figure 3D, see also Table S3). Unsupervised clustering revealed that Inhibitor TBI (SD cMet) and Inhibitor TBI (SD VEGFR) groups were comparable with each other, but were significantly different from Saline TBI and Saline sham groups. On the other hand, the Inhibitor TBI (SD Btk) group was quite similar to the saline TBI group. Differential expression analysis revealed 137 targets with phosphorylation status significantly affected by single dose of cMet/HGFR or VEGFR inhibitors, with only 19 targets shared by the two treatments (Figure 3E). Most importantly, treatment with single dose of cMet/HGFR inhibitor caused the down-phosphorylation of the microglial-specific signaling proteins Btk and Plcg2 as well as of RTKs involved in microglial activation and microgliosis such as c-Kit and IL-3R (Bright et al., 2004; Spiller et al., 2018). Interestingly, the cMet/HGFR inhibitor caused significant down phosphorylation of hepatocyte growth factor regulated tyrosine kinase substrate (HRS), responsible for promoting degradation of activated RTK (Gao and Woude, 2015), and increase in phosphorylation levels of the adapter Blnk, reversing the activation pattern observed in saline TBI group. The Inhibitor TBI (SD cMet) also showed broader impact, affecting the signaling of EGFR, and VEGFR2. Interestingly, VEGFR2 phosphorylation levels were significantly upregulated in 3H low impact, compared to 3H high impact but similar to Inhibitor TBI (SD cMet) group (cfr. Table S1, Figure 3E). Comparatively, Inhibitor TBI (SD VEGFR) significantly increased the phosphorylation levels of CDK5 and IGF1R responsible for microglial activation (Muyllaert et al., 2008; O’Donnell et al., 2002), and downregulation of immune-related IL7R/CD127, protein associated with resting microglial phenotype (Saijo and Glass, 2011) (Table S3).

**Figure 3.**
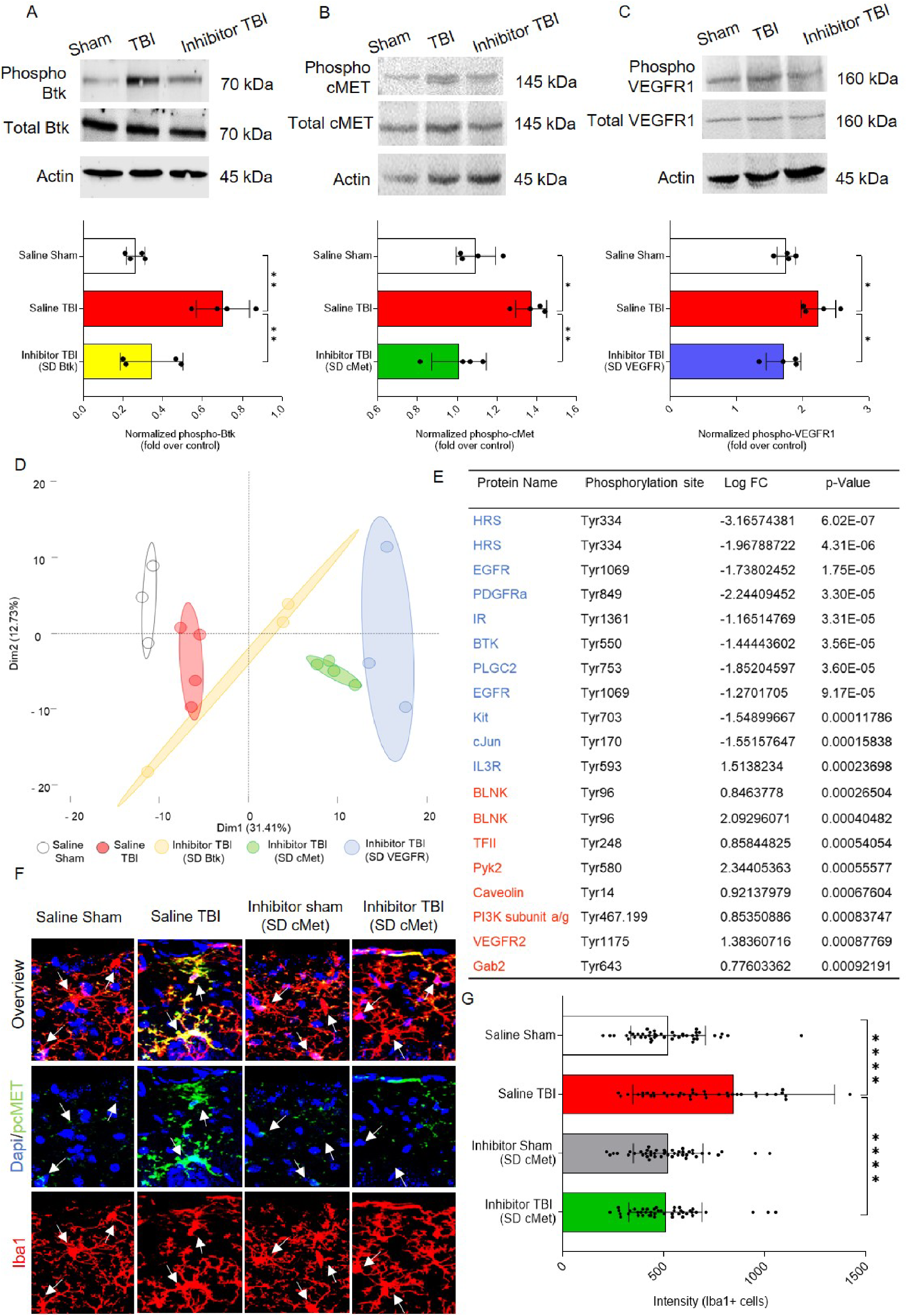
Inhibitor treatment alters the signaling pattern post trauma. (A)-(C). At 3h after TBI, levels of (A) phosphorylated Btk (Tyr222), (B) phosphorylated cMet/HGFR (Tyr1235), and (C) phosphorylated VEGFR1 (Tyr1333) were significantly upregulated in saline TBI samples. Respective inhibitor treatment significantly reduced phosphorylated levels to baseline. (See also Figure S3 for full scan of blots). (D) Post preprocessing principal component analysis (PCA) plot showing group-based clustering with minimal overlap among groups. (E) List of DE proteins (p<0.05) up-phosphorylated (red) or down-phosphorylated (blue) for comparisons between Saline TBI and Inhibitor TBI (pcMet) with logarithmic fold change and individual significance. (F)-(G) Immunostaining shows significant reduction in phospho-cMet/HGFR expression levels upon inhibitor treatment. The graph shows computed intensity levels of phosphorylated cMet/HGFR expression among four groups. All graphs are represented as mean ± SD. In (A)-(C), n=4 per group. Dots indicate individual animals. Significance of differences between means were analyzed using one-way anova test with Tukey correction (recommended) (*P<0.05, **P<0.01). In (D)-(E), n=3 for Saline sham, Inhibitor TBI (SD Btk), Inhibitor TBI (SD VEGFR1) and n=4 for Saline TBI and Inhibitor TBI (SD cMet) groups. Significance for DE proteins was set at P<0.05 (BH adjusted). Detailed individual comparisons between Saline TBI and Inhibitor TBI (SD VEGFR1) for the proteins are shown in Table S3. In (F)-(G), n=4 per group. In (G), dots indicate intensity of individual cells. Significance of differences between means were analyzed using two-way anova test with Tukey correction (recommended) (****P<0.0001).

### Blockade of cMet/HGFR prevents microglial activation and limits vascular and neuronal stress induced by TBI

The combination of tyrosine phosphorylation landscape analysis, phospho-epitope immunostaining and the effect of small-molecule inhibitors on pTyr signaling pointed to a previously unappreciated role of cMet/HGFR in regulating microglial responses to TBI.

As a prerequisite of further exploration of cMet/HGFR regulation of microglial activation, we verified that the single dose of cMet/HGFR inhibitor indeed significantly decreased the phospho-cMet/HGFR immunoreactivity in microglia in cortical sections obtained 3h after TBI (Figure 3F, G; Saline TBI vs saline sham: p=0.0001, saline TBI vs inhibitor TBI (SD cMet): p=0.0045).

Next, we conducted a morphometric analysis of microglial cells obtained from sham-operated mice and in mice subject to TBI after saline or SD cMet/HGFR inhibitor administration (Figure 4A). Whereas microglia in TBI samples displayed a retraction of processes compared to sham samples (a marker of microglial activation; Fernandez-Arjona et al., 2019, Morrison et al., 2017), treatment with the single dose of cMet/HGFR inhibitor preserved and actually enhanced the overall size of microglial arborization, a morphology closely resembling that of resting microglia (Figure 4B, C; branch length: Saline TBI vs saline sham: p=0.045, saline TBI vs inhibitor TBI (SD cMet): p<0.0001, branch length and endpoints: saline TBI vs inhibitor TBI (SD cMet): p<0.0001).

**Figure 4.**
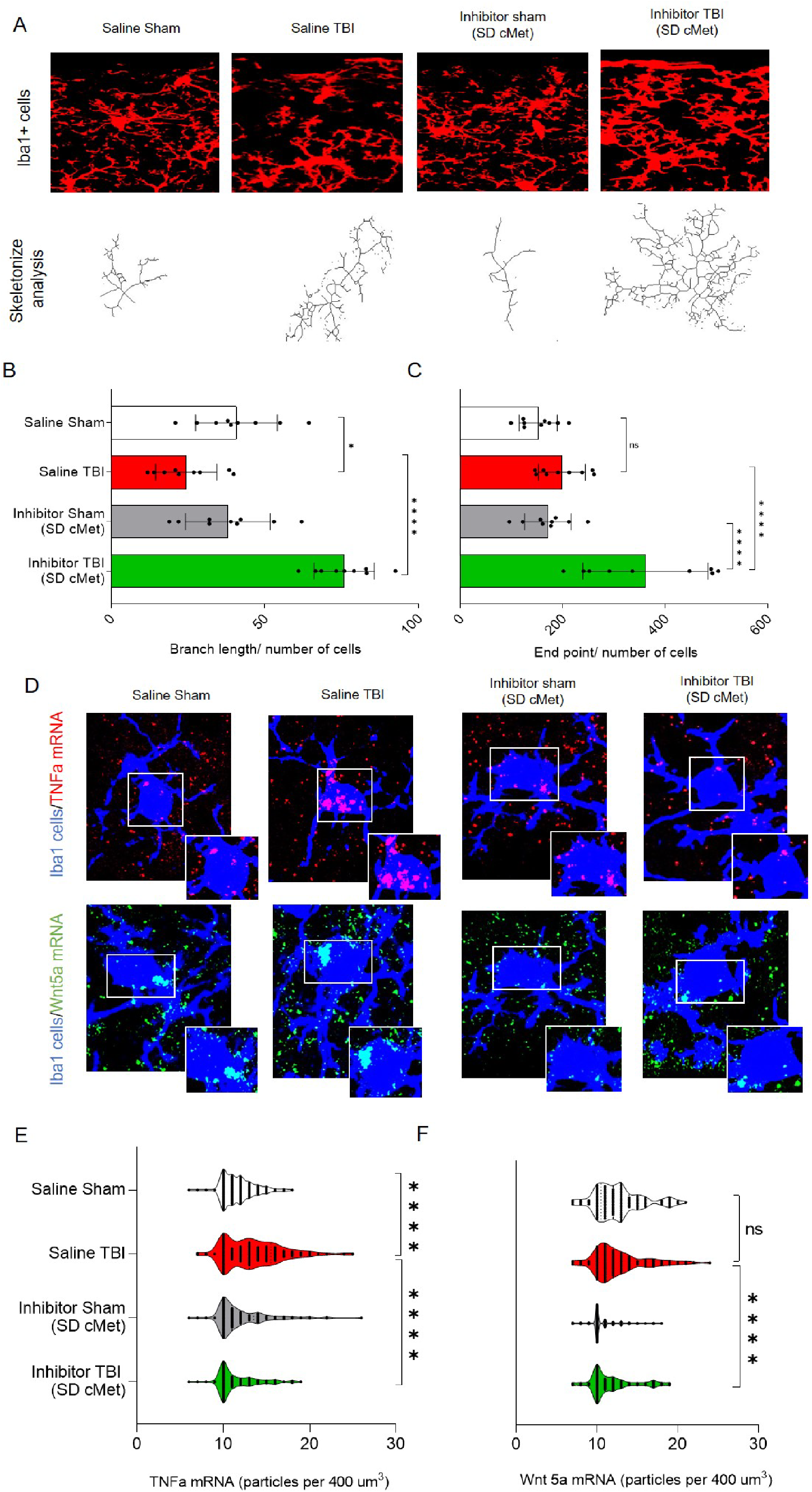
Impact of acute cMet/HGFR blockade on morphological changes and alteration in inflammatory mediators in microglia. (A) Acute morphological changes in microglia showing decrease in branch length within 3h post trauma in the saline treated group depicting activated microglia. (B-C) The graph plots show that upon single dose (SD) cMet inhibitor treatment, the (B) branch length and (C) endpoints were significantly increased displaying a ramified or arborized microglia morphology. (D) At 3h post TBI, In situ hybridization showing TNF-α mRNA level (Red) in microglia was significantly upregulated upon TBI and decreased to baseline with inhibitor treatment. In comparison, Wnt-5a mRNA (Green) was not affected upon TBI, but was significantly downregulated post inhibitor treatment. (E)-(F) The volcano plot shows quantification for the mRNA levels of (E) TNF-α and (F) Wnt-5a, statistically supporting the data. In (A)-(C), n=3 per group. Dots indicate individual sections per animal. Significance of differences between means were analyzed using two-way anova test with Tukey correction (recommended)(ns=not significant, *P<0.05, ****P<0.0001). In (D)-(F), n=4 per group dots indicate mRNA count per cell. Significance of differences between means were analyzed using two-way anova test with Tukey correction. (ns=not significant, ****P<0.0001).

Next, we assessed if blockade of cMet/HGFR would affect the induction of microglial inflammatory mediators, in particular the induction of TNF-α, among the first events unfolding upon TBI (Fan et al., 1996; Frugier et al., 2010). Single-molecule in situ hybridization revealed that TNF-α mRNA was almost undetectable in microglia at baseline but was significantly upregulated upon TBI; notably, such upregulation was negated by the pretreatment with the single dose of cMet/HGFR inhibitor (Figure 4D, E; Saline TBI vs saline sham: p<0.0001, Saline TBI vs Inhibitor TBI (SD cMet): p<0.0001). We explored a second modulator of microglial function, namely Wnt5a. Wnt5a is a paracrine mediator highly expressed in so-called M1 inflammatory microglia (Mecha et al., 2020) and able to induce chemotaxis and iNOS, COX2 expression in microglia (Halleskog et al., 2012) and in other phagocytes (Yu et al., 2014). Interestingly, Wnt5a mRNA was not significantly upregulated by TBI in microglia, but was dramatically suppressed by the cMet/HGFR inhibitor (Figure 4D, F; Saline TBI vs Inhibitor TBI (SD cMet): p<0.0001), further stressing the profound immunomodulatory role of this receptor.

Finally, we aimed at dissecting the impact of acute cMet/HGFR blockade on broad TBI responses. Since our model has not been fully characterized on these terms, we first determined the proteome changes occurring in the cortex at 3h. We detected a total of 6059 proteins among which 36 were significantly upregulated and 25 were significantly downregulated (compared to sham samples; Figure 5A, B. See also Table S4). It is worth noting that some of the proteins with the highest fold-change (>2 fold) are involved in epigenetic responses and RNA metabolism (Hells, Syf2, Prpf38b), cytoskeletal remodeling (Pak4 and Pak2, Plnxa3), several enzymes and ion transporters (Ppcs, Scl12a6, Scl4a7), neuro-vascular inflammation (Itgb3), the matrix protein Vitronectin (Vtn), several transport proteins (Kif5a, Zwilch) and vasculature-related proteins (Vwa5a).

**Figure 5.**
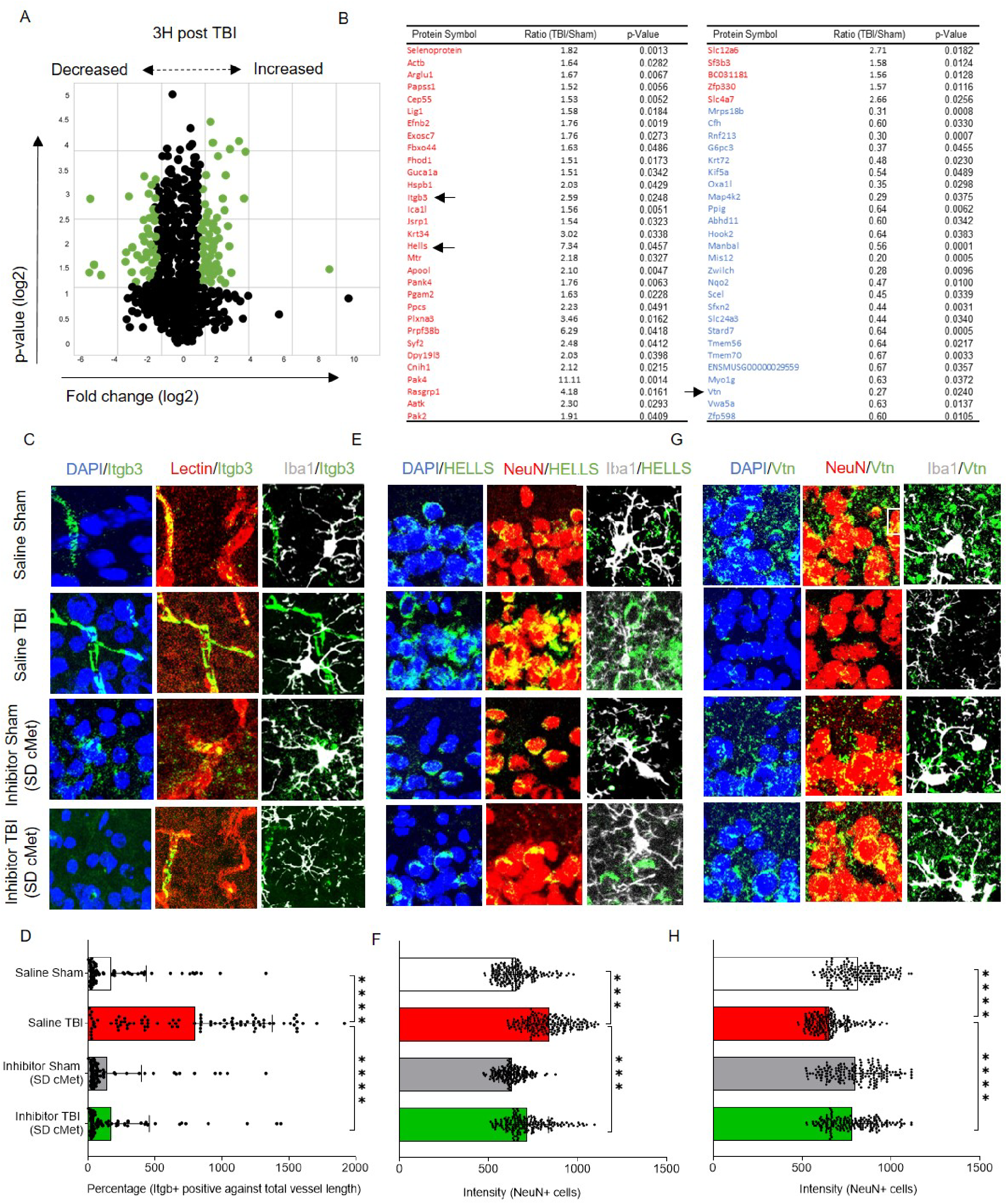
Broad range impact of the cMet/HGFR inhibitor on proteomic alteration. (A) Post proteome analysis volcano plot showing distribution of proteins of the injured cortex compared to sham at 3h post injury. After applying a cutoff value for fold change (1.5 times above or below the normalised values) and p-value (p<0.05, FDR corrected), significant proteins were selected. (B) List of statistically significant upregulated (Red) and downregulated (Blue) proteins with fold change and individual significance. Table S4 provides details including the function of significant proteins. (C)-(H) Selected proteins were immunostained to verify proteomic changes 3h post TBI and identify inhibitor effect. Figure S4 displays overview images for the zoomed insets. (C) Integrin beta 3 (itgb3) colocalized with vascular marker (Lectin) and showed no colocalization with neurons (NeuN) or microglia (Iba1). (D) The bar graph shows the percentage of itgb3 levels (Itgb3+ area against total vessel length) were significantly upregulated to approximately 2.5 folds, 3h post trauma and reduced to baseline levels with single dose cMet (SD cMet) inhibitor treatment. (E) Lymphoid specific helicase (HELLS) colocalized with neurons (NeuN) and showed no overlapping with microglia (Iba1+) cells. (F) The bar graph shows the intensity levels of HELLS were significantly upregulated 3h post trauma whereas reduced to baseline levels with single dose cMet (SD cMet)inhibitor treatment. (G) Vitronectin (Vtn) colocalized with neurons (NeuN) and showed no overlapping with microglia (Iba1+) cells (H) Vtn levels were significantly downregulated after TBI and single dose cMet (SD cMet) inhibitor treatment restored Vtn levels to baseline levels. All graphs are represented as mean ± SD. In (A)-(B), n=3 per group. Significance for DE proteins was set at P<0.05 (FDR adjusted). In (C)-(H), n=4 per group. In (D), (F) and (H), dots indicate individual cells/vessels. Significance of differences between means were analyzed using two-way anova test with Tukey correction. (***P<0.001, ****P<0.0001). In (D), expression levels were normalized over the total length of the vessel.

We considered three of these proteins (upon further validation) to monitor the impact of the single dose of cMet/HGFR inhibitor, namely Integrin beta-3 (Itgb3 an adhesion molecule expressed on microglia and on endothelial cells and involved in leukocyte extravasation), Lymphocyte-Specific Helicase (HELLS; involved in epigenetic regulation and RNA processing in neurons [Lai et al., 2020]) and the Vitronectin (an extracellular matrix protein involved in neuronal and microglial physiology; Welser-Alves and Milner, 2013; Del Zoppo et al., 2007]).

As before, mice were treated with a single dose of cMet/HGFR inhibitor (or saline) and subject to TBI or sham surgery before being sacrificed at 3 h post injury. The levels of Itgb3, Hells and Vtn were assessed by immunostaining and their expression pattern was verified against markers of neuronal, microglial and vascular subpopulations (labelled by NeuN, Iba1+ and lectin, respectively, Figure 5C-H. See also Figure S4). Igbt3 was expressed at a very low level in microglia in sham-operated mice, but displayed a massive upregulation in endothelial cells upon TBI (Figure 5C, D; Saline TBI vs saline sham: 5-fold increase, p<0.0001). Notably, vascular Itgb3 upregulation was completely abolished by the single dose of cMet/HGFR inhibitor (Figure 5C, D; p<0.0001). Hells was expressed only in neurons; its levels were significantly upregulated after injury (p<0.0001) but not when cMet/HGFR was blocked (Figure 5E, F; p<0.0001). Vtn immunoreactivity was diffused, with clusters localized in proximity of neurons (NeuN+; Figure 5G). A substantial loss of Vtn levels upon TBI (p=0.0001). However, cMet/HGFR inhibition prevented this effect (Figure 5G, H; p=0.015); Taken together, these data demonstrate that interference with cMet/HGFR activation in microglial cells produce a normalizing effects on TBI-induced responses, including vascular inflammation, neuronal stress and matrix remodeling.

### Acute and prolonged cMet/HGFR inhibitor treatment prevents acute TBI-induced motor impairment

We hypothesized that blockade of acute microglial reaction by the cMet/HGFR inhibitor would result in beneficial outcomes at behavioral level. Since our TBI model generates a focal lesion in correspondence of the somatosensory cortex, we exploited a quantitative test involving sensorimotor integration, based on a single pellet reaching task. Animals were trained for 10 days to retrieve food pellets from a narrow slit using their forelimbs; success rates were recorded before the TBI and at 1, 3, 5, 7, 10, 14, 17 and 21 dpi (Figure 6A). For each animal, success and failure rates were assessed before and after TBI (as % of total attempts/trials). We quantified: (i) success rate (successful retrieval of the pellet), (ii) failure in reaching (highlighting impairment in motor adaptation during reaching), (iii) failure in grasping (underscoring the impairment in digit movements), (iv) failure in retrieval (underscoring digit control impairment). Animals subjected to trauma were pretreated with a single dose of cMet/HGFR inhibitor followed by daily vehicle for 7 days (SD cMet n=4), multiple doses (MD) of cMet/HGFR inhibitor starting from 2h before TBI and followed by a daily administration of the inhibitor for 7 days (MD cMet; n=6) or only saline (Saline TBI; n=7). For comparison, we also administered a single dose of Btk inhibitor (SD Btk; n=3) and a single dose of VEGFR inhibitor (SD VEGFR; n=4) to independent groups (Figure 6A). In saline-treated animals (Saline TBI group), the success rate significantly decreased as early as 1 dpi compared to pretreatment recordings (Figure 6B; Red: p<0.0001 vs pre-trauma). This was largely due to an increase in reaching failure affecting the spatial perception post trauma (Pavlides et al., 1993, Vidoni et al., 2010, Mathis et al., 2017), whereas grasping and retrieval failures remained consistently similar (Figure 6C; Red: p<0.0001 vs pre-trauma, Figure 6D). A single dose of cMet/HGFR inhibitor; Inhibitor TBI (SD cMet) group, (administered 2h before TBI) resulted in a significant preservation of success rate at 1 dpi (Figure 6B; Green: p=0.0005 vs 1dpi Saline TBI group) that remained statistically better than the saline-treated groups also at 2 dpi (p=0.0023 vs Saline TBI group) and 3 dpi (p=0.0492 vs Saline TBI group). The improved success rate was mirrored by the selective decrease in reaching failure, indicating a preserved functioning of the somatosensory network (Figure 6C; Green: p=0.048 vs 1dpi Saline TBI group). A single dose of the VEGFR inhibitor; Inhibitor TBI (SD VEGFR), also resulted in a preservation of the performance at 1dpi (Figure 6B; Blue: p=0.048 vs 1dpi Saline TBI group), but immediately lost protective effects from 2dpi . A single dose of the Btk inhibitor; Inhibitor TBI (SD Btk), failed to produce any significant preservation of success rate or improvement in failure rate (Figure 6B,C; Black).

**Figure 6.**
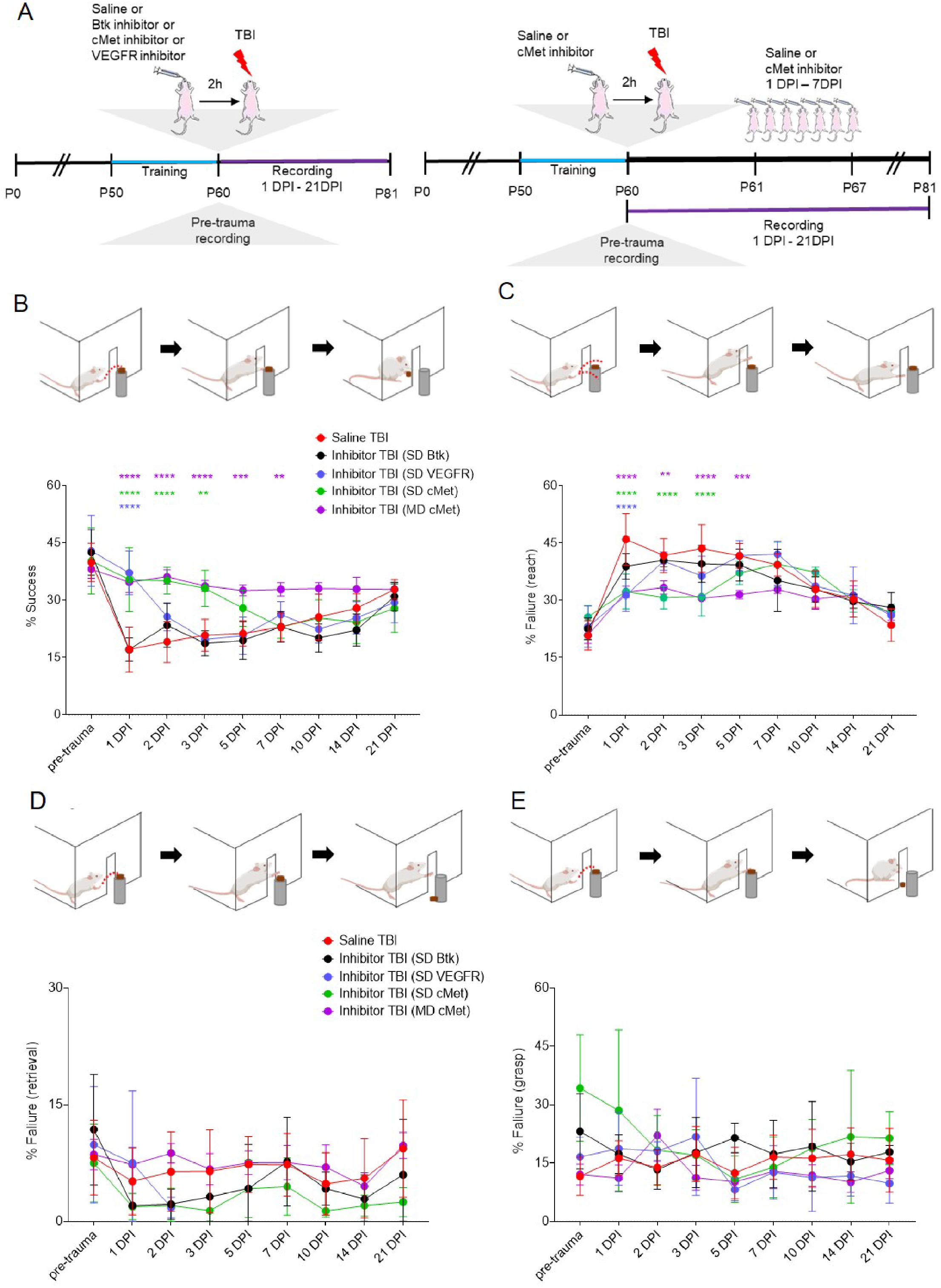
cMet/HGFR inhibitor elicits beneficial effects on TBI-induced motor impairment. (A) Schematic outline of the experiment with 5 groups; saline TBI, inhibitor TBI (SD Btk), inhibitor TBI (SD VEGFR1), inhibitor TBI (SD cMet) and inhibitor TBI (MD cMet) to investigate treatment effect. Single dose (SD) administered 2h before trauma or multiple doses (MD), starting 2h before trauma for consecutive 7 days were recorded pre and post trauma until 21 days. (B) The graph shows success rate significantly reduced at 1 DPI until 14 DPI (saline TBI group; Red) as well as in inhibitor TBI (SD Btk; Black) compared to respective pre-treatment groups. Whereas the success percentage of two groups; inhibitor TBI (SD cMet; Green) and inhibitor TBI (SD VEGFR; Blue), was significantly improved compared to Saline TBI group with a single dose of inhibitor. Inhibitor TBI (SD cMet) group showed a persistent improvement in success percentage until 3 DPI compared to Saline TBI group. Success percentage was significantly improved with consecutive multiple doses of cMet treatment [inhibitor TBI (MD cMet); purple] starting 1 DPI with persistent improvement until 7 DPI. (C) The graph shows a significant increase in reaching failure (highlighting impairment in motor adaptation during reaching) in Saline TBI and Inhibitor TBI (SD Btk) group at 1DPI. A significant decrease in reaching failure was observed in Inhibitor TBI (SD cMet) and inhibitor TBI (SD VEGFR1) groups compared to Saline TBI group. Inhibitor TBI (SD cMet) group showed a persistent decrease in reaching failure until 3 DPI compared to Saline TBI group. Inhibitor TBI (MD cMet) group shows significant decrease in the percentage of reaching failure compared to Saline TBI group from 1 DPI until 5 DPI. (D-E) The graph shows failures in (D) retrieval (underscoring digit control impairment) and (E) grasping (underscoring the impairment in digit movements) among five treatment groups. No significant changes or patterns were observed among the groups. All graphs are represented as mean ± SD. In (A)-(C), n=7 for Saline TBI, n=3 for inhibitor TBI (SD Btk), n=4 for inhibitor TBI (SD VEGFR1) and inhibitor TBI (SD cMet), and n=6 for inhibitor TBI (SD cMet) treatment. Significance of differences between means were analyzed using two-way anova test with Tukey correction. (**P<0.001,***P<0.001, ****P<0.0001).

Most notably, mice that received the cMet/HGFR inhibitor for 7 consecutive days; Inhibitor TBI (MD cMet), (administered 2h before trauma and after behavioral testing for successive days), displayed a persistent preservation of the success rate compared to Saline TBI group (Figure 6B: Purple; p<0.0001 vs 1dpi Saline TBI, p<0.0001 vs 2dpi Saline TBI, p=0.0002 vs 3dpi Saline TBI, p=0.0012 vs 5dpi, p=0.0099 vs 7dpi saline TBI) and a significantly reduced reaching failure rate during the treatment period (between day 1 and day 7, Figure 6C: Purple; p<0.0001 vs 1dpi Saline TBI, p=0.0131 vs 2dpi Saline TBI, p<0.0001 vs 3dpi Saine TBI, p=0.012 vs 5dpi Saline TBI). The improvement in success rate was fully attributed to a reduced reaching failure (Figure 6C,D,E). Taken together these data suggest that TBI impairs motor performance by reducing spatial perception while cMet/HGFR inhibitor treatment preserves the functioning of somatosensory networks.

### Preserved neuronal integrity and resting microglial morphology upon cMet/HGFR inhibition in TBI

Finally, we explore if the beneficial effect of single dose or multiple doses of cMet/HGFR inhibitor on sensorimotor performance would have a counterpart on the overall neuronal loss and post-traumatic gliosis. Mice were treated with either a single dose or multiple doses with a 7-days course of the cMet/HGFR inhibitor or with saline and were sacrificed at 21 dpi (Figure 6A). In saline treated mice (Saline TBI), the density of NeuN+ cells was significantly reduced in the site of injury (Figure 7A, B; p<0.0001 vs saline sham). Notably, a single dose of cMet/HGFR inhibitor [Inhibitor TBI (SD cMet)] resulted in a significant increase in the number of surviving neurons in the injury site (Figure 7A, B; p<0.0001 vs Saline TBI). Likewise, mice administered with cMet/HGFR inhibitors for 7 days [Inhibitor TBI (MD cMet)] displayed a significantly larger number of surviving NeuN+ cells in the injury area compared to saline treated TBI mice, although still lower than in sham mice (Figure 7A, B p<0.0001 vs Saline TBI).

**Figure 7.**
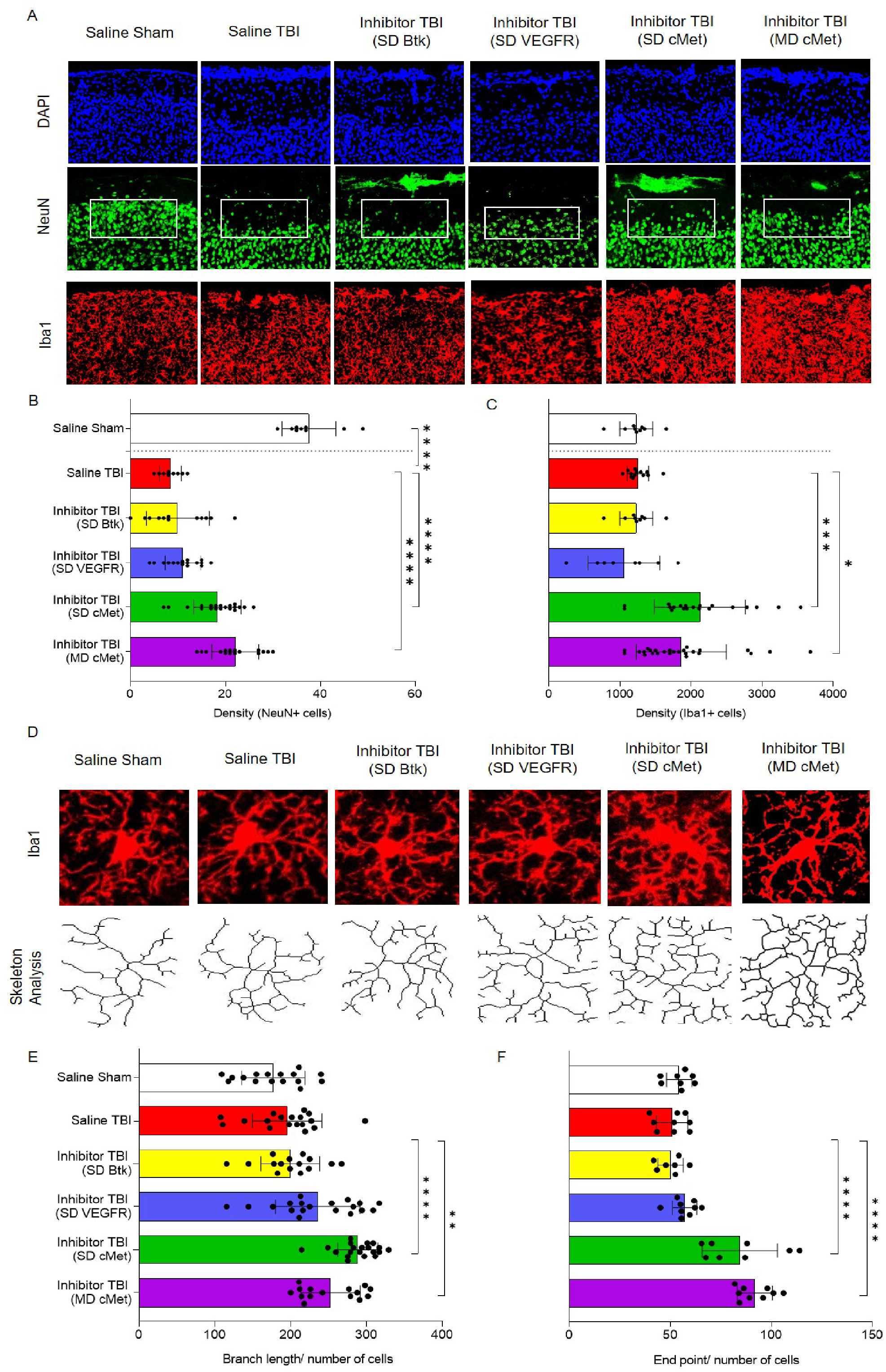
Acute cMET/HGFR inhibitor treatment preserves neuronal density and branched microglial morphology. (A) Immunostaining showing decreased number of neurons in saline TBI group compared to saline sham and increased number of neurons in Inhibitor TBI (SD cMet) and inhibitor TBI (MD cMet) treatment group compared to saline TBI group. Microglia cells also show increase in density among Inhibitor TBI (SD cMet) and inhibitor TBI (MD cMet) groups. (B) The bar graph shows significant decrease in the neuronal density in saline TBI group compared to saline sham and significant improvement in neuronal density upon single dose and multiple doses of cMet inhibitor. (C) The bar graph shows significant increase in microglial density computed over the area as well as single cells upon single dose or multiple doses of cMet inhibitor. (D) Morphological changes in microglia showing increase in branch length and endpoints per cell upon single or multiple doses of cMet inhibitor treatment displaying a ramified or arborized microglia morphology. (E)-(F) The bar graph shows significant increase in (E) branch length per cell and (F) endpoints per cell upon cMet inhibitor treatment compared to saline TBI group. All graphs are represented as mean ± SD. In (A)-(F), n=7 for Saline TBI, n=3 for inhibitor TBI (SD Btk), n=4 for inhibitor TBI (SD VEGFR1) and inhibitor TBI (SD cMet), and n=6 for inhibitor TBI (SD cMet) treatment. Dots represent cells/recording per individual sections. Significance of differences between means were analyzed using two-way anova test with Tukey correction. (*P<0.05, **P<0.001,***P<0.001, ****P<0.0001).

We further analyzed the density and morphology of microglial cells at this stage. Single dose of cMet/HGFR inhibitor [Inhibitor TBI (SD cMet)] displayed significant increase in overall microglial density (a composite readout including cell number and cell size; Figure 7A,C, p<0.0001 vs Saline TBI) however the same effect was not observed for Inhibitor TBI (SD Btk) or Inhibitor TBI (SD VEGFR) group. We further analyzed the morphometrical changes of microglia in the different groups. The results revealed no standing difference in saline TBI group compared to saline sham controls, but a statistically significant increase in branch length and number of end points per cell in both cMet/HGFR inhibitor treated groups: single dose (Figure 7D-E; Inhibitor TBI (SD cMet); Branch length: p<0.0001 vs Saline TBI, Endpoint; p<0.027 vs Saline TBI) as well as multiple doses (Figure 7D-F; Inhibitor TBI (MD cMet); p<0.01 vs Saline TBI). Single dose of Btk inhibitor [Inhibitor TBI(SD Btk)] or VEGFR inhibitor [Inhibitor TBI(SD VEGFR)] displayed no increase in the branch length or the number of endpoints per cell with no significant long-term changes in microglial morphology. Taken together the results suggest cMet/HGFR inhibitor treatment results in an improved protection of neuronal integrity and increases the preservation of a non-reactive microglial morphology, identified as a ramified resting state, even 21 dpi after injury.

### HGF levels are acutely upregulated in the CSF of human TBI patients

Our murine data collectively demonstrate the role of HGF/cMet signaling in driving the acute reactivity of microglia upon TBI. We sought to validate the relevance of these findings to human subjects by assessing the levels of HGF in the Cerebrospinal Fluid (CSF) obtained from human TBI patients (clinical and demographic data in Table 1). Compared to non-traumatic CSF controls, HGF levels were very strongly increased already at D0, partially declined at D1 and returned to values closer to the baseline by D3 (Figure 8A). In a subset of patients that was longitudinally sampled a similar trend was observed, with massive elevation in the early hours and quick decline toward the baseline. In order to compare the time course of HGF, we measured the levels of two other pro-inflammatory cytokines, IL-6 and IL-1β, known to increase in acute (IL-6) or sub-acute (IL-1β) phases (Chiaretti et al., 2005; Kumar et al., 2015; Yan et al., 2014). Levels of IL-6 were very elevated at D0, and quickly declined over the next three days; on the other hand, IL-1β showed a positive trend increasing from D0 to D3. Thus, HGF trend was similar to that of IL-6 and differed from IL-1β; in fact, HGF levels were strongly correlated with IL-6 levels in each sample. Thus, HGF upregulation characterizes the acute phases of human TBI, in agreement with what is seen in murine samples.

**Figure 8.**
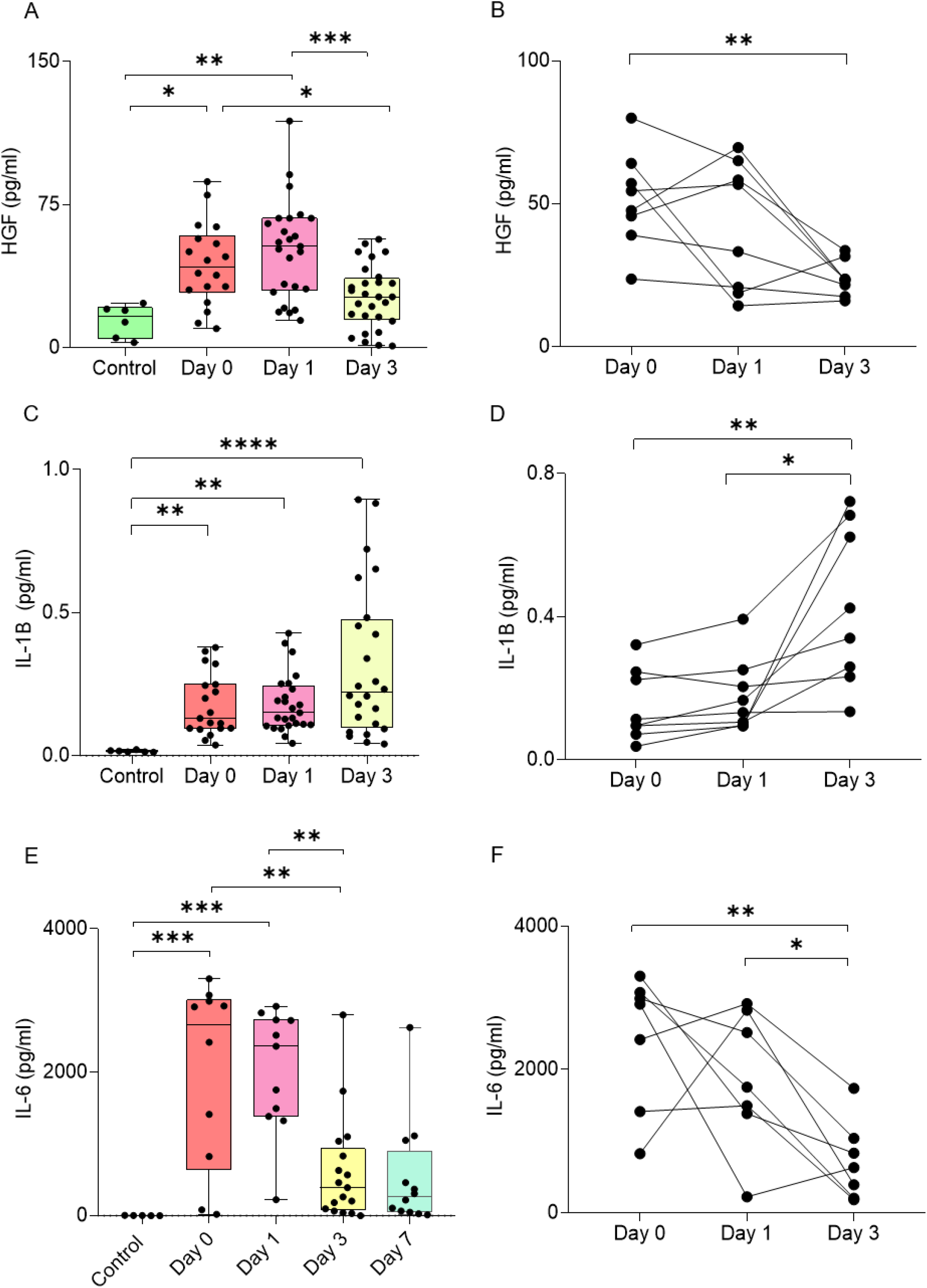
HGF is expressed in the human brain and is upregulated upon TBI in the human CSF. Significant upregulation of (A-B) HGF, (C-D) IL-1B, and (E-F) IL-6 protein in cerebrospinal fluid samples from traumatic brain injury patients compared to control samples. ELISA results for (B) HGF and SIMOA results for (F) IL-6 show upregulation at 0 and 1 day after trauma and reduce after 3 days. On the other hand, (D) IL-6 levels increase over time from day 0 to day 3. All graphs are represented as mean ± SD. In (A), (C), (E), n=6 for control, n=18 for day 0, n=25 for day 1, n=28 for day 3, n=12 for day 7. In (B), n=8 for all groups. In (D), (E), n=7 for all groups. Dots represent individual human samples. Data sets were checked for normality. Significance of differences between means were analyzed using two-way anova test with Tukey correction. (*P<0.05, **P<0.001,***P<0.001, ****P<0.0001)

**Table 1.** Clinical and demographic data for control subjects and TBI patients

|  | **Control group\*** | **D0 group** | **longitudinal group** |
| --- | --- | --- | --- |
| N (M/F) | 6 (2/4) | 28 (21/7) | 11 (7/4) |
| Age (median, range) | 71 (53-85) | 34.5 (21-55) | 29 (23-42) |
| Glasgow Coma Scale (median, range) | N/A | 6 (3-9) | 6 (3-7) |
| Injury Severity Score (median, range) | N/A | 34.5 (17-59) | 43 (17-59) |
| Glasgow Outcome Scale-Extended (median, range) | N/A | 4 (1-8) | 4 (1-6) |

\* non-TBI CSF samples were obtained from hydrocephalus patients (without previous TBI) undergoing implantation of ventriculoperitoneal shunts.

## Discussion

TBI is characterized by the sudden and simultaneous unfolding of neuronal, glial and vascular responses to injury, which involves multiple signaling cascades setting in motion; cell death, local inflammation, synapse disruption, glial activation and blood brain barrier disruption. We have captured the complexity of these events initially by monitoring the phosphorylation status of 39 different RTK at multiple timepoints after TBI. This approach has the advantage to focus on targets with high translational potential (Tyrosine kinases) while identifying the signature of several concomitant processes and cell subtypes.

Our findings confirm previous anecdotal reports about individual RTKs (Chandrasekar et al., 2018; Ding et al., 2014; Kyyriäinen et al., 2017) and further extend the characterization of the acute signaling landscape in TBI. Notably, the 3h/1dpi samples display changes in the phosphorylation of a set of receptor involved in microglial reactivity, namely the upregulation of the phosphorylation of pro-inflammatory cMet/HGFR and MSPR, together with the concomitant downregulation in the phosphorylation of reactive microglia-suppressing (Ray et al., 2017; Fourgeaud et al., 2016) Axl and Dtk (Figure 1C,D), pointing toward a coordinated shift in the balance between different signals regulating microglial activation. Notably, this effect is not present for milder TBI (where Axl and Dtk are actually up-phosphorylated), indicating that even small differences in TBI severity may translate in either the enhancement or the suppression of the reactive microglial phenotype (Figure S1I).

Further characterization of the signaling landscape at 3h using a distinct array platform and immunohistochemistry confirmations, not only confirmed several of the targets already identified (most notably cMet/HGFR, VEGFR1 and EGFR, underscoring the robustness of the findings) but revealed the phosphorylation of additional RTK/NRTK involved in microglia reactivity and neuroinflammation, such as Flt3 (Anandasabapathy et al., 2011) and a series of NRTK such as Fer, Lck, Abl and Btk. The latter is largely restricted to immune cells, including microglia, and it is involved in microglial reaction to LPS (Nam et al., 2018) and stroke (Ito et al., 2015). We elected to use small-molecule inhibitors to explore the roles of cMet/HGFR, VEGFR and Btk on microglial activation and on the overall secondary damage. Small molecules provide the advantage to deliver acute loss-of-function experiments, free of adaptations that constitutive gene knock-out may produce and offer a more direct translational outlook. Both the cMet/HGFR inhibitor and the VEGFR inhibitor exerted a substantial impact on the phosphoproteome: the cMet/HGFR inhibitor caused the down-phosphorylation of a number of RTK involved in immune function (c-Kit, IL3R, PDGFRa) and of related signaling molecules, most notably Btk itself, Blnk and of Hrs (the Hepatocyte growth factor-Regulated tyrosine kinase Substrate), a critical signaling hub for cMet/HGFR (Abella et al., 2005; Row et al., 2005). The VEGFR inhibitor affected a much larger set of targets, coherently with the broader expression of the receptor itself (and the broader specificity of the inhibitor). Interestingly, the VEGFR-inhibitor affected targets partially overlapped with the cMet/HGFR one, in particular with the down-phosphorylation of Btk, Hrs, IL3R and EGFR. The Btk inhibitor itself does not appear to be sufficient to bring about a significant change in the phosphoproteome or a beneficial effect in histological and behavioural readouts. This finding is in contrast with the previously established beneficial effect of Btk suppression in stroke (Ito et al., 2015) and in demyelinating disease (Pellerin et al., 2021). Indeed, the limited effect of the Btk inhibitor on the signaling cascades, despite an impact on Btk phosphorylation itself, suggest that either the redundancy of the signaling cascades is so that inhibition of a single downstream/non-receptor kinase is less effective than the inhibition of upstream receptors (which lead to Btk down-phosphorylation) or that a more intense suppression of Btk is needed to observe beneficial effects.

We have mechanistically investigated the role of cMet/HGFR blockade on microglial activation (and its broad consequences in the acute TBI landscape). We demonstrated that the assumption of a reactive morphology and the induction of inflammatory mediators in microglia upon TBI is prevented by cMet/HGFR blockade, and that vascular inflammation, matrix degradation and neuronal stress are also prevented. Finally, the grasping task revealed that either a single dose or a 7-days multiple dose treatment with cMet/HGFR inhibitor delivered a significant improvement in motor performance, corresponding to an enhanced preservation of neuronal integrity in the site of injury.

Taken together, our dataset identifies cMet/HGFR as a new regulator of microglial activation and, more broadly, of the neuroimmunological response to TBI (Figure 9). cMet/HGFR and the closely related RON receptor are expressed on microglia (Cohen et al., 2014; Di Renzo et al., 1993; Suzuki et al., 2008; Yamagata et al., 1995), macrophages (Nishikoba et al., 2020; Zhao et al., 2015) and other immune cells including subsets of B and T lymphocytes (Benkhoucha et al., 2020; Gordin et al., 2010) as well as in a number of non-neuronal tissues (Organ and Tsao, 2011). In macrophages, cMet/HGFR induces motility, chemotaxis and chemokine secretion (Beilmann et al., 2000; Zhao et al., 2015), and proliferation (Moransard et al., 2010). Interestingly, cMet/HGFR has been reported to shift macrophage polarization toward the so-called non-inflammatory M2 phenotype (Choi et al., 2019; Nishikoba et al., 2020) and stimulates IL-10 secretion (Coudriet et al., 2010). On the other hand, cMet/HGFR activation enhances macrophages infiltration of joints and chemokine secretion (Beilmann et al., 2000; Hosonuma et al., 2021) and drives the proliferation of M1-polarized macrophages (Moransard et al., 2010). Thus, cMet/HGFR appears to mediate recruitment and activation of macrophages, and contribute to polarization of the phenotype, although with opposite effects in different microenvironments. While microglia expresses both cMet/HGFR and its ligand HGF (Yamagata et al., 1995), the consequences of cMet/HGFR signaling in microglia in a pathology context are unclear. Our data suggest that cMet/HGFR activation may gate the induction of a reactive phenotype and contribute to the induction of inflammatory cytokines. Additional mechanisms may be involved in the overall effect of cMET/HGFR inhibition on microglia. Given the role in inducing chemotaxis, cMet/HGFR may contribute (together with other chemotactic factors, e.g., purinergic receptors; Roth et al., 2014) to microglial migration toward the site of injury. Interestingly, microglial cells remained in a ramified-morphology state up to 14 days after the last administration of the inhibitor, suggesting that cMet/HGFR may gate the induction of the reactive phenotype in the acute phase and, when inhibited, permanently prevents access to this phenotype later on.

**Figure 9.**
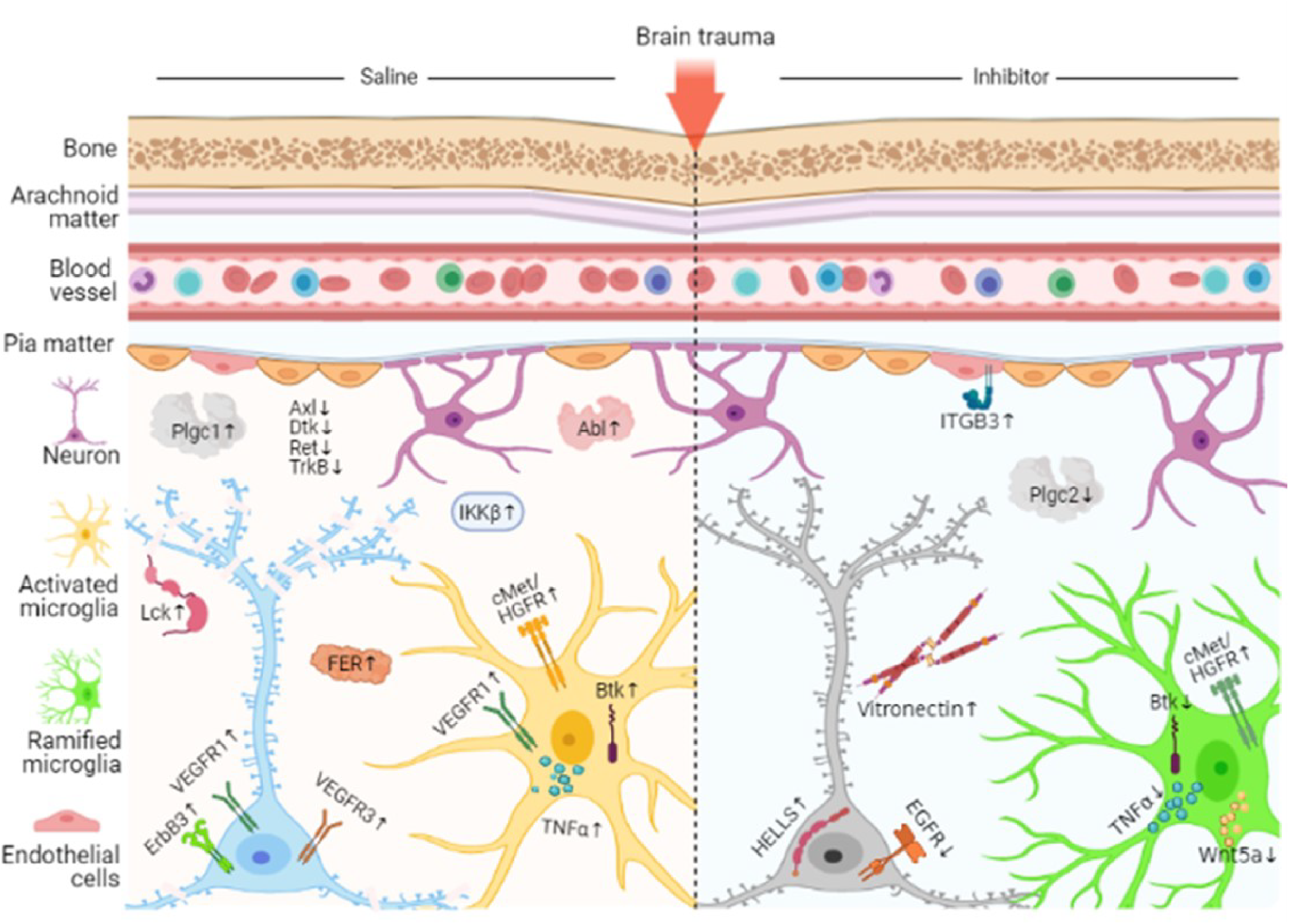
Temporal mapping of tyrosine kinase phosphorylation and related proteins upon trauma and alterations upon cMet/HGFR inhibitor treatment at 3H post injury. Trauma alters the phosphorylation pattern of tyrosine kinase receptors by increasing the levels of pcMet/HGFR, pVEGFR1, pVEGFR3, pErbB3, pBTK as well as decreasing levels of pAxl, pDtk, pcRet and pTrkB, among several other downstream signaling proteins. Inhibition of phosphorylated cMet/HGFR disrupts the signaling and affects broad range targets resulting in increased arborization and ramified microglial morphology leading to neuroprotection and improved behavioral outcome.

The present study is not without limitations. First, our exploration of the role of cMet/HGFR, VEGFR and Btk relies on small-molecule inhibitors. This approach has the advantages to explore selectively acute effects, devoid of long-term or developmental adaptations due to constitutive gene knock-out and, at the same time, to provide insights toward drug repurposing strategy (therefore delivering a better translational outlook). On the other hand, small-molecule TK inhibitors are rarely completely selective, and inhibition of closely related RTK (e.g., simultaneous blockade of VEGFR1, VEGFR2 and VEGFR3) may contribute to the overall effect. Moreover, the efficacy of small-molecule inhibitors is affected by their pharmacokinetics, in particular by their blood-brain-barrier penetration. Therefore, weak effects may not necessarily point to irrelevance of the target but may be confounded by insufficient target engagement. Second, we elected to investigate a TBI model with relatively mild (in terms of NSS score and histology) consequences and the findings may be not directly extrapolated to conditions characterized by large hematomas, extensive necrotic lesions or bone fracture. In fact, even a small difference in kinetic energy may drive distinctive responses (as shown in this paper) and hematomas may drive substantially different biology (Chandrasekar et al., 2018). However, the elevation of HGF seen in a clinically representative cohort of human subjects suggests that our experimental findings may be applied to human subjects.

Nevertheless, our findings identify a number of RTK with transient activation in the early phases of the TBI responses; the cMet/HGFR inhibition shows that interventions at this stage may have profound acute as well as long-lasting modulatory effects on microglial response to TBI, neuronal integrity and motor performance. Thus, cMet/HGFR provides a proof-of-concept for the identification of new modulators of TBI response based on their impact on the acute large-scale signaling landscape which may shape the overall response to trauma in the long term.

## Author contribution

FR conceived the study and supervised the project. RR and FR planned the experiments. RR and MEW performed R software-based analyses. RR and MM performed the behavior experiments. RR, MM and LE analyzed the behavioral data. RR, MG and JK performed and analyzed the proteomic experiments. AC and MMK collected the human CSF samples. RR and SSK performed the analysis of the human CSF. RR, FoH and AC performed the trauma surgery and collected the samples for histological analysis. RR prepared figures and drafted the manuscript. FR, AT, MG, TB, AL, MMK and RR authors contributed to the final version of the manuscript. All authors read and approved the manuscript.

## Materials and methods

### Resource table

| REAGENT or RESOURCE | SOURCE | IDENTIFIER |
| --- | --- | --- |
| **Antibodies** |  |  |
| Mouse anti-NeuN | Abcam | PA5-78499 |
| Rabbit anti-GFAP | Abcam | Ab4674 |
| Chicken anti-GFAP | Abcam | Ab4674 |
| Rabbit anti-pMet/HGFR<br>(Tyr1234) | Cell signaling | 3077S |
| Rabbit anti-pMet/HGFR<br>(Tyr1234) | R&D systems | AF2480 |
| Rabbit anti-pBTK (Tyr222) | Novus biologicals | NBP1-78295 |
| Rabbit anti-pVEGFR1<br>(Tyr1333) | Biorbyt | orb644609 |
| Rabbit anti-HELLS | Antibodies.com | A34212 |
| Rat anti-Vitronectin | R&D systems | MAB38751 |
| Mouse anti-Integrin beta 3 | Novus biologicals | NB600-1342 |
| Guinea pig anti-Iba1 | Synaptic systems | 234004 |
| Lycopersicon Esculentum<br>(Tomato) Lectin (LEL, TL),<br>DyLight® 594 | Vector labs | DL-1177-1 |
| **Chemicals and Inhibitors** |  |  |
| Alexa Fluor 488 Tyramide SuperBoost Kit | Invitrogen | B40922 |
| JNJ-38877605 | Selleckchem | HY50683 |
| Spebrutinib (CC-292) | Selleckchem | S7173 |
| Valatanib (PTK-787) | MedChemExpress | HY-12018 |
| **Critical Commercial Assays** |  |  |
| Proteome RTK array |  |  |
| Full Moon Tyrosine phosphorylation array | Fullmoon Biosystems | PCS300 |
| Antibody array assay kit | Fullmoon Biosystems | KAS02 |
| **Software and Algorithm** |  |  |
| Image J | NIH | NA |
| Prism 7.0 | GraphPad Software | NA |
| R studio | R | NA |
| GenePix Pro Software v7 | Molecular Devices LLC | 5004757 |
| **Other** |  |  |
| LEICA Confocal microscope | DMi8 S Platform | NA |
| GenePix 4000B Microarray Scanner | Molecular Devices LLC | GENEPIX 4000B-U |

### Mouse strains

All experimental procedures were performed in compliance with animal protocols approved by the local veterinary and animal experimentation committee at University Ulm, Germany under the license number 1370. B6SJL male mice aged between p60–p90 days were used throughout the study.

### Antibodies

Primary antibodies used were: Mouse anti-NeuN (1:100, Abcam), Rabbit anti-GFAP (1:500, Abcam), Rabbit anti-pMet/HGFR (1:200, Cell Signaling/R&D systems), Rabbit anti-pBTK (1:200, Novus Biologicals), Rabbit anti-pVEGFR1 (1:200, Biorbyt), Rabbit anti-HELLS (1:100, Antibodies.com), Rat anti-Vitronectin (1:100, R&D systems), Mouse anti-Integrin beta 3 (1:50, Novus biologicals), Guinea pig anti-Iba1 (1:500, Synaptic systems), Chicken anti-GFAP (1:500, Abcam) and Lycopersicon Esculentum (Tomato) Lectin (LEL, TL), DyLight® 594 (1:100, Vector labs). Secondary antibodies were used from Invitrogen or Life Technologies, raised in either goat or donkey against primary antibody’s host species, highly crossed absorbed and conjugated to fluorophores of Alexa Fluor 488, Alexa Fluor 568, Alexa Fluor 647 and used at a 1:500 dilution. Alexa Fluor 405 conjugated DAPI was used at 1:1000 dilution. For signal amplification, streptavidin conjugated to Alexa Fluor 488 (1:1000, Life Technologies) was used to amplify against the biotin tag.

### Traumatic brain injury

For all surgical procedures, mice were anesthetized with sevoflurane/isoflurane (2-4% in 96% O2) and were subcutaneously injected with buprenorphine (0.1mg/kg) as a pre-or postoperative analgesic. TBI was induced in mice by a modified closed, blunt weight drop model (previously reported Flierl et al., 2009). The scalp was shaved and eye ointment was applied post-operatively to protect the cornea. Scalp skin was then incised on the midline to expose the skull and the animals were positioned in the weight-drop apparatus in which the head was secured to a holding frame. Using the 3-axis mobile platform in the apparatus, the impactor was positioned to the coordinates of the injection site (From bregma ≈ x = +3.0mm, y = − 2.0mm, z = 0.0mm).

TBI was delivered by a weight of 120/125 g dropping from a height of 45 cm (or 40 cm for low impact). A mechanical stop prevented a skull displacement (by the impactor) larger than 2.5 mm, in order to limit brain damage. Apnea time was monitored after injury. The overall time of mice exposure to sevoflurane/isoflurane did not exceed 10 mins. After restoration of normal breathing, mice were removed from the stereotactic frame. The skin was sutured with Prolene surgical thread and the mice were left on the heating pad until regained consciousness. Animals were then transferred to recovery cage ad libitum access to food and water. Additional buprenorphine doses were administered every 12 hours until 3 days. To avoid unnecessary suffering, mice were checked every 2 hours on day 1 and then every day until 7 days post injury for TBI determining fixed termination criterion. Effort was made to minimize the suffering of animals and reduce the number of animals used.

### Perfusions and tissue processing

Animals were given an overdose of anesthesia and transcardially perfused with 25 mL ice cold PBS 0.5M EDTA followed by 50mL 4% paraformaldehyde (PFA) [company name]. After perfusion, brains were dissected and post fixed in 4% PFA overnight at 4 . Tissues were cryo protected by sinking in 30% sucrose in PBS. Samples were frozen in the Optimal Cutting Temperature (OCT) compound (Tissue Tek, Sakura) using dry ice. 40 μ thick free-floating sections, spanning the injection/TBI site (identified using coordinates, appearance of third and lateral ventricles, corpus callosum using Allen brain Atlas as reference) were cut in a cryostat, collected in PBS and processed for immunostaining.

### Protein extraction

For proteomic analysis, mice were sacrificed at respective timepoints (or sham surgery in case of control) by placing them in heavy sevoflurane filled containers. Once the animals stopped breathing, they were removed from the container and the brain was quickly dissected in cold PBS. Tissue samples (Ø 0.25 cm) were punched from the lesioned and contralateral uninjured somatosensory cortex and immediately frozen at −80°C to extract proteins, samples were then thawed in RIPA buffer prepared in house (150mM NaCl, 10mM Tris, 0.1% Sodium Dodecyl Sulfate (SDS), 1% Triton 100X and 5mM EDTA) containing Phosphatase (1 tablet per 3.3ml lysis buffer) and Protease (1 tablet per 16.67ml lysis buffer) inhibitor (Roche cOmplete tablets, Sigma-Aldrich, Taufkirchen, Germany) cocktail and homogenized with approximately 20 strokes of Dounce apparatus. Tissue homogenates were then cleared by centrifugation twice (10.000g, 10min and 10.000g, 5 mins) and assayed for protein concentration using the BSA kit from ThermoFisher.

### Phospho RTK array processing

Proteome Profiler Mouse Phospho-RTK Array Kit (R&D Systems, Minneapolis) was used to determine the Phospho RTK activation pattern. The nitrocellulose membrane arrays provided in the kit were based on sandwich immunoassay and processed according to manufacturer’s instructions. Briefly, membranes spotted with the anti-RTK antibody were blocked in Array buffer 1 for 1h at RT. 130 ug of extracted protein was diluted in 1.5mL Array buffer 1 overnight at 4°C. The membranes were then washed 3 x 10 min in Wash Buffer and incubated for 2 hours at RT with Anti-Phospho-Tyrosine-HRP Detection Antibody, diluted to 1:5000 in 1X Array Buffer 2. After final washing steps, HRP detection was performed by adding 1 mL Clarity Max™ Western ECL Blotting Substrates from Bio-Rad. Arrays were imaged using BioRad X-ray imager and quantified using ImageJ. ROI was drawn on each antibody spot with a constant diameter and mean gray value was recorded. Further analysis was performed using R software.

### Tyrosine Kinase Array processing

After determining the protein concentration in the protein lysate, 65 ug of protein was loaded to the arrays. The arrays were processed according to the manufacturer’s instructions. Briefly, the sample was biotinylated with Biotin/DMF solution for 2 hours at RT with episodic vortexing. The solution was mixed with reaction stop reagent and incubated for 30 mins at RT. The glass arrays were incubated in a blocking solution for 45 minutes and washed 10 times with distilled water until the slide was smooth without any residue. After adding a biotinylated sample to the coupling solution, arrays were incubated for 2 hours at RT and washed with 1X wash buffer three times for 10 mins. The arrays were washed again with distilled water (G biosciences) and incubated with a detection buffer with Cy5-streptavidin (ThermoFisher) at 1:1000 for 20 minutes at RT protected from light. The washing steps were repeated as described above. After removing excess water from the slides, the arrays were dried using 10 psi compressed air at a 30 degree angle with 2 inch distance between the nozzle and the glass slide. The arrays were images using GenePix 4000B array scanner (Molecular Devices, LLC) and the image analysis was performed using GenePix Pro Software v7 (Molecular Devices, LLC). The settings for the analysis were kept constant in all cases. The GAL file was loaded in the software and the ROI was adjusted on the protein spots. Each intensity on F635 was recorded and GPR files were saved. Further analysis was performed using R software.

### Array analysis

Chemiluminescence signal for each spot was logged after microarray image analysis. The raw intensity values for each receptor/protein were recorded automatedly via image recorder software or manually using ImageJ software. The raw data files were loaded in R software and the dataset for each array was preliminary subjected to quality control assessment (QCA); outlier identification, data distribution, intra-array and inter-array normalization. Normalized data for each array was subjected to principal component analysis (PCA) to display group-based clustering. Confidence ellipses (assuming multivariate normal distribution) with the first two principal components were plotted to validate further analysis. Modified linear modeling-based analysis was then applied to the data to identify RTK showing a significant increase or decrease in phosphorylation at the different timepoints. For protein array analyses, the code has been made publicly available on open-access GitHub repository PROTEAS (PROTein array Expression AnalysiS; github.com/Rida-Rehman/PROTEAS).

### Pharmacological agents and treatment procedures

The BTK inhibitor CC-292 (Selleckchem chemicals, Munich, Germany), cMet/HGFR inhibitor JNJ38877605 (Medchem) and VEGFR1 inhibitor Vatalanib (PTK787) 2HCl (Selleckchem chemicals, Munich, Germany) are commercially available. Drugs were dissolved in a minimal volume of DMSO; CC-292 10mg in 50ul, JNJ38877605, 10mg in 150 ul, PTK787, 10mg in 100ul. The selected doses for CC-292: 30mg/kg (Evans et al. 2013, Levin et al. 2015, Tonge et al. 2018), JNJ38877605: 40mg/kg (Etnyre et al. 2014, Lolkema et al. 2015, De Bacco et al. 2016), and PTK 787: 50mg/kg (Reardon et al. 2009, Shankar et al. 2016, Kong et al. 2017) were dissolved in 200 ul of vehicle (90% saline, 5% PEG 400 and 5% Tween 80) maintaining the concentration of DMSO less than 5% in the final dose. The inhibitors were orally administered at doses 2 hours before the trauma to ensure maximum occupancy in the brain until procedure is performed.

For longer inhibitor treatment in behavior experiments, drug or saline was administered 2h before trauma for a period of 7 consecutive days. For each successive day, dose was administered immediately after the behavior session.

### Immunostaining

Before blocking, antigen retrieval was performed by incubating the sections in Sodium Citrate Buffer (10mM Sodium Citrate, 0.05% Tween 20, pH 6.0) in a hot water bath (75L) for 45 minutes. For phosphorylated proteins, sections were then subjected to hydrogen peroxide (0.3% H_2_O_2_) treatment for 20-30 minutes to quench the endogenous peroxidase activity. Sections were then blocked in a blocking buffer (3% BSA, 0.3% Triton in PBS) for 2 h at 24 °C on a rotary shaker. Following blocking an appropriate mix of primary antibodies were diluted in the blocking buffer and incubated for 48 h at 4 °C. For staining against phosphorylated proteins, sections were then incubated in poly-HRP-conjugated secondary antibody or HRP-conjugated streptavidin from Alexa Fluor 488 Tyramide SuperBoost Kit (Invitrogen) for 60 minutes and then incubated in Tyramide working solution according to manufacturer’s protocol. After incubating in Reaction stop reagent for 8 mins, sections were washed in PBS for 3 × 30 min and incubated in the appropriate mix of other secondary antibodies, together with, whenever appropriate, the DNA dye TOPRO-3 (1:1000, Invitrogen), diluted in a blocking buffer for 2 h at 24 °C. After further washing the sections for 3 × 30 min in PBS, the sections were mounted on coverslips and onto the slides using FluoroGold Plus (Invitrogen).

### Microscopy

Confocal images were acquired using an LEICA DMi8 S inverted microscope, fitted with a 10× air or 20× oil objective. Tile-scans of 5×3 were acquired to cover the full span of the injury site and the full cortical thickness. The images were acquired in a 12-bit format. Imaging parameters (laser power, photomultiplier voltage, digital gain, and offset) were established with the goal of preventing saturation in target structures while obtaining a lowest signal intensity of at least 150 (in arbitrary units). Image capture and processing conditions were kept constant when imaging was used for quantification. Brightness and contrast of images are adjusted and pseudo-colored for presentation.

### Image analysis

Region of interest was 2×10µm^2^ and kept constant for all analyses. For the measurement of NeuN+ cells, Iba1+ cells and GFAP+ cells, 5 × 3 composite tiles cans of confocal stacks were acquired with the 10 × objective to image the injury site and the surrounding penumbral and perilesional areas. Confocal stacks of 10–12 optical sections (at the same depth) were collapsed in maximum-intensity projections in ImageJ and a threshold was set for the resulting images, to establish a reproducible criterion. Regions of Interests (ROIs) were positioned centered on the axis of the injury site. In these ROIs, the number of NeuN+ cells and/or GFAP+ cells were manually counted. At least 3–5 tissue sections were analyzed from each of 3–6 mice per experimental group.

For p-cMet/HGFR, pBTK, pVEGFR1, Vitronectin and HELLS fluorescence intensity analysis, confocal stacks composed of 19-21 optical sections (acquired at the same depth in the tissue section) were collapsed in maximum-intensity projections using the ImageJ software and a threshold was set for the resulting images, to establish a reproducible criterion. Microglia and neurons were identified based on their morphology, size and positive immunostaining for the microglial marker (Iba1) and neuronal marker (NeuN), respectively. A constant target region was considered for each experiment, spanning the cortical layer II–III and centered on the axis of the injury site. ROIs encompassing the cellular soma (excluding the nucleus for neurons) were manually drawn for each cell in the target region and the integrated average fluorescence intensity was logged. A minimum of 100 cells (layer II–III) from 3 distinct tissue sections from each of 3–5 mice were quantified. The median fluorescence intensity of phosphorylated receptors was computed. Microglia morphology was quantified using imagej skeleton and fractal analysis procedures as mentioned in (Morrison et al., 2018).

For Integrin beta 3 intensity analysis, confocal stacks composed of 19-21 optical sections (acquired at the same depth in the tissue section) were collapsed in maximum-intensity projections using the ImageJ software and a threshold was set for the resulting images, to establish a reproducible criterion. A target region was considered, spanning the cortical layer II– III and centered on the axis of the injury site. Vessels were stained using a vascular marker (Collagen IV) and area was recorded for a minimum 20 vessels of variable lengths in each section. After applying intensity threshold, area positive for Integrin beta 3 across these vessel lengths were recorded and normalised to compute the percentage of positive area.

### Behavioral performance and assessment

The overall neurological/motor impairment was evaluated using forelimb reaching task recorded using four high-speed cameras (200 frames/second) for subsequent analysis. Animals were trained for 10 days for 20 minutes to extract food pellets from a narrow slit using their forelimbs. Food was restricted 2 hours after the training session until the next session or conclusion of the experiment. Exclusions were made based on a 10% reduction in body weight of animals. After the training period, pre-treatment recordings were made. Post trauma recordings were made after 1, 3, 5, 7, 10, 14, 17 and 21 days. The analysis was performed by manually counting four modular aspects of forelimb movement. The trial was considered only if the paw is extended outside the plexiglass slit; (i) success; retrieving the pellet inside the container, (ii) failure (reach); unable to reach food due to problem in spatial understanding, (iii) failure (grasp); reaches the pellet but unable to grasp it properly and (iv) failure (retrieval); hold the pellet but unable to retrieve inside the container. Percentages for the recorded trials for all four conditions were recorded by averaging against respective total trials per animal per session and analysis were performed by comparing groups with pretreatment (for TBI group) or subsequent time points among different groups.

### In situ Hybridization

Single-molecule in situ mRNA hybridization was performed according to manufacturer instructions (ACDBio, RNAscope, Fluorescence In Situ mRNA Hybridization for histological sections, all reagents/buffers were provided by ACDbio) with some modifications. Briefly, slides were mounted on glass slides. Target retrieval was performed for 3 min and sections were washed with dH20 and ethanol. Thereafter, slides were pretreated with protease III for 20 min at 40 °C and probe hybridization (for Wnt5a and TNFa) was performed for 4.5 h at 40 °C, followed by 2 × 2 min washing step with washing buffer (provided by ACDbio) and 30 min incubation of amplification-1 reagent at 40 °C. Slides were washed 2 × 2 min and incubated with amplification-2 reagent for 15 min at 40 °C followed by another 2 × 2 min washing step. The last amplification was performed by 30 min incubation with amplification-3 followed by a 2 × 2 min washing step. A final detection amplification was performed by incubating the amplification-4 reagent for 45 min at 40 °C and the final washing step was increased to 2 × 10 min. The co-immunostaining procedure was performed soon after as follows: slides were blocked for 1 h in 10% BSA, 0,3% Triton in PBS and incubated overnight with primary antibody (Guinea pig anti-Iba1, 1:500 [abcam] and mouse anti-NeuN, 1:500 [abcam]). Slides were washed 3 × 30 min and incubated with secondary antibodies (Donkey anti-mouse 405, 1:500 [invitrogen] and Donkey anti-guinea pig 647 [Invitrogen]). After the last round of washing (3 × 30 min in PBS), the slides were mounted using Fluorogold prolong antifade mounting medium (Invitrogen)

### Human CSF samples

CSF samples from TBI patients were obtained at the Alfred Hospital, Melbourne and was approved by the Alfred ethical committee and by the ethical committee of Ulm University (licence 194/05). Informed consent for the sampling of CSF was obtained from the next of the kin; data regarding the original cohort have been previously published (Yan et al., 2014).

Inclusion criteria were: severe TBI with a post-resuscitation GCS ≤8 (unless initial GCS>9 was followed by deterioration requiring intubation) and, upon CT imaging, the need for an extraventricular drain (EVD) for ICP monitoring and therapeutic drainage of CSF. Exclusion criteria comprised pregnancy, neurodegenerative diseases, HIV and other chronic infection/inflammatory diseases, or history of TBI. CSF was collected over 24 h and kept at 4°C; samples were obtained on admission (day 0), in the first 24h (day 1) and three days after injury (day 3). CSF samples were centrifuged at 2000g for 15 min at 4°C and stored at -80°C until analysis. Out of the 42 TBI patients constituting the original cohort (Yan et al., 2014), we selected samples from 30 patients, depending on the availability of three aliquots for day 0, day 1 and day 3 after injury (Full details available in Table 1).

### ELISA and SIMOA assays

Single-molecule arrays were performed as described as per manufacturer instructions (Quanterix) for the quantification of IL-6 and IL-1β CSF samples were diluted 1:20 for IL-6 measurements and diluted 1:5 for IL-1β measurements. The assay was then read on HD-1 Analyzer (Quanterix, USA). HGF levels in CSF were measured by sandwich ELISA for HGF (DuoSet Kit, R&D Systems, Minneapolis, MN, USA), as described by the manufacturer. The detection range was 125-8000 pg/mL for HGF. Undiluted CSF samples were used and the assay was read on a microplate reader set to 450 nm wavelength.

### Quantitative and statistical analysis

Statistical analysis was performed with the GraphPad Prism software suite. Mann-Whitney U-Test was used for two group comparisons, One-way ANOVA with Bonferroni Correction was performed among three groups and Two-way ANOVA with Tukey correction was used for four group comparisons to examine statistical significance. Protein array analysis was performed using R software. Error bars represent standard deviation (SD), unless indicated otherwise. Statistical significance was set at P < 0.05.

## Supplementary figure legends

**Figure S1. Changes in cellular survival/infiltration and protein phosphorylation over time post TBI**

(A) Position of Region of Interest (ROI) in the imaged brain section to display injury core and surrounding penumbra in layer II/III.

(B) Immunolabeling of neurons (NeuN, green) and astrocytes (GFAP, red) at sham, 3HPI, 1DPI, 3DPI and 7DPI to ascertain the impact elicited by weight-drop trauma model.

(C)-(D) The histogram displays the number of (C) neurons and (D) astrocytes significantly increasing at 3DPI and peaking at 7DPI.

(E)-(H) Graphs displaying the expression pattern of individual proteins over time.

(I) Comparison of RTK phosphorylation pattern between two different intensities of trauma.

All graphs are represented as mean ± SD. In (C)-(D), n=4 for sham, 3HPI, 3DPI and 7DPI. Dots, squares and triangles indicate the number of cells per section and four sections per animal. Significance of differences between means were analyzed using one-way anova test with bonferroni correction. (*P<0.05, **P<0.05, ***P<0.0001). In (E)-(I), n=6 for sham, 1DPI, 3HPI low intensity, 3DPI and 7DPI and n=7 for 3HPI high intensity. Significance for DE proteins was set at P<0.05 (FDR adjusted). Scale bars: 200µm and 300µm for (A) and (B), respectively. Detailed individual comparisons between respective groups and the overlapping phosphorylation pattern for the proteins are shown in Table S1.

**Figure S2. Cell specific expression of selected tyrosine kinases**

(A)-(C) Immunostaining was performed to identify neuronal expression of (A) phosphorylated cMet/HGFR, (B) Phosphorylated Btk and (C) phosphorylated VEGFR1. No significant difference in p-cMet/HGFR and pBtk levels was observed post trauma however pVEGFR1 levels were elevated after trauma.

(D) Vascular co-staining showed no overlap between p-cMet/HGFR, pBtk, but slight overlap in pVEGFR1 expression.

**Figure S3. Full scans of western blots data**

**Figure S4. Overview images for the zoomed insets of selected proteins**

## Supplementary tables

**Table S1.** Detailed information of significant proteins (Related to Figure 1 and Figure S1). This table details the independent interaction between two groups to evaluate the phosphorylation pattern of RTK families.

**Table S2.** Detailed information of significant proteins (Related to Figure 2). This table covers the significant proteins identified during large scale in-depth screening of tyrosine kinase proteins at 3H post trauma.

**Table S3.** Detailed information of significant proteins (Related to Figure 3). This table covers the significant proteins altered upon acute VEGFR inhibitor treatment.

**Table S4.** Detailed information of significant proteins (Related to Figure 5). This table covers the significant proteins identified from analyzing the proteome changes occurring in the cortex at 3H post trauma.

## Acknowledgement

The present work has been supported by the ERANET-NEURON initiative. FR, MG, AT, MM, JK and RR are members of the MICRONET consortium (funded by BMBF: FKZ 01EW1705A for FR). MG is independently funded by the TRR274. MM is supported by FWO as part of the MICRONET consortium in the ERANET-NEURON initiative grant awarded to AT (G0H1817N). RR is also independently funded by the Hannelore Kohl Foundation Award. FR is also funded by DFG in the context of the SFB1149 (DFG No. 251293561) as well as through individual grants (DFG No. 443642953, 431995586 and 446067541).

## References

1. Abella, J. V, Peschard, P., Naujokas, M.A., Lin, T., Saucier, C., Urbé, S., and Park, M. (2005). Met/Hepatocyte Growth Factor Receptor Ubiquitination Suppresses Transformation and Is Required for Hrs Phosphorylation. Mol. Cell. Biol. 25, 9632–9645.

2. Alam, A., Thelin, E.P., Tajsic, T., Khan, D.Z., Khellaf, A., Patani, R., and Helmy, A. (2020). Cellular infiltration in traumatic brain injury. J. Neuroinflammation 17, 328.

3. Anandasabapathy, N., Victora, G.D., Meredith, M., Feder, R., Dong, B., Kluger, C., Yao, K., Dustin, M.L., Nussenzweig, M.C., Steinman, R.M., et al. (2011). Flt3L controls the development of radiosensitive dendritic cells in the meninges and choroid plexus of the steady-state mouse brain. J. Exp. Med. 208, 1695–1705.

4. Bai, L., Bai, G., Wang, S., Yang, X., Gan, S., Jia, X., … & Yan, Z. (2020). Strategic white matter injury associated with longLterm information processing speed deficits in mild traumatic brain injury. Human brain mapping, 41(15), 4431–4441.

5. De Bacco, F., D’Ambrosio, A., Casanova, E., Orzan, F., Neggia, R., Albano, R., Verginelli, F., Cominelli, M., Poliani, P.L., Luraghi, P., et al. (2016). MET inhibition overcomes radiation resistance of glioblastoma stemLlike cells. EMBO Mol. Med. 8, 550–568.

6. Beilmann, M., Vande Woude, G.F., Dienes, H.P., and Schirmacher, P. (2000). Hepatocyte growth factor-stimulated invasiveness of monocytes. Blood 95, 3964–3969.

7. Benkhoucha, M., Senoner, I., and Lalive, P.H. (2020). C-Met is expressed by highly autoreactive encephalitogenic CD8+ cells. J. Neuroinflammation 17, 68.

8. Bright, J.J., Natarajan, C., Sriram, S., and Muthian, G. (2004). Signaling through JAK2-STAT5 Pathway Is Essential for IL-3-Induced Activation of Microglia. Glia 45, 188–196.

9. Chandrasekar, A., Olde Heuvel, F., Wepler, M., Rehman, R., Palmer, A., Catanese, A., Linkus, B., Ludolph, A., Boeckers, T., Huber-Lang, M., et al. (2018). The Neuroprotective Effect of Ethanol Intoxication in Traumatic Brain Injury Is Associated with the Suppression of ERBB Signaling in Parvalbumin-Positive Interneurons. J. Neurotrauma 35, 2718–2735.

10. Chiaretti A, Genovese O, Aloe L, Antonelli A, Piastra M, Polidori G, Di Rocco C. Interleukin 1beta and interleukin 6 relationship with paediatric head trauma severity and outcome. Childs Nerv Syst. 2005 Mar;21(3):185–93; discussion 194. doi: 10.1007/s00381-004-1032-1. Epub 2004 Sep 29. PMID: 15455248

11. Choi, W., Lee, J., Lee, J., Lee, S.H., and Kim, S. (2019). Hepatocyte growth factor regulates macrophage transition to the M2 phenotype and promotes murine skeletal muscle regeneration. Front. Physiol. 10.

12. Cohen, M.E., Fainstein, N., Lavon, I., and Ben-Hur, T. (2014). Signaling through three chemokine receptors triggers the migration of transplanted neural precursor cells in a model of multiple sclerosis. Stem Cell Res. 13, 227–239.

13. Coudriet, G.M., He, J., Trucco, M., Mars, W.M., and Piganelli, J.D. (2010). Hepatocyte growth factor modulates interleukin-6 production in bone marrow derived macrophages: Implications for inflammatory mediated diseases. PLoS One 5.

14. Dash, P.K., Moore, A.N., and Dixon, C.E. (1995). Spatial memory deficits, increased phosphorylation of the transcription factor CREB, and induction of the AP-1 complex following experimental brain injury. J. Neurosci. 15, 2030–2039.

15. Davalos, D., Grutzendler, J., Yang, G., Kim, J. V., Zuo, Y., Jung, S., Littman, D.R., Dustin, M.L., and Gan, W.B. (2005). ATP mediates rapid microglial response to local brain injury in vivo. Nat. Neurosci. 8, 752–758.

16. Del Zoppo, G.J., Milner, R., Mabuchi, T., Hung, S., Wang, X., Berg, G.I., and Koziol, J.A. (2007). Microglial activation and matrix protease generation during focal cerebral ischemia. In Stroke, pp. 646–651.

17. Dey, A., Allen, J.N., Fraser, J.W., Snyder, L.M., Tian, Y., Zhang, L., Paulson, R.F., Patterson, A., Cantorna, M.T., and Hankey-Giblin, P.A. (2018). Neuroprotective role of the ron receptor tyrosine kinase underlying central nervous system inflammation in health and disease. Front. Immunol. 9.

18. Ding, K., Wang, H., Xu, J., Lu, X., Zhang, L., and Zhu, L. (2014). Melatonin reduced microglial activation and alleviated neuroinflammation induced neuron degeneration in experimental traumatic brain injury: Possible involvement of mTOR pathway. Neurochem. Int. 76, 23–31.

19. Elmore, M.R.P., Najafi, A.R., Koike, M.A., Dagher, N.N., Spangenberg, E.E., Rice, R.A., Kitazawa, M., Matusow, B., Nguyen, H., West, B.L., et al. (2014). Colony-stimulating factor 1 receptor signaling is necessary for microglia viability, unmasking a microglia progenitor cell in the adult brain. Neuron 82, 380–397.

20. Etnyre, D., Stone, A.L., Fong, J.T., Jacobs, R.J., Uppada, S.B., Botting, G.M., Rajanna, S., Moravec, D.N., Shambannagari, M.R., Crees, Z., et al. (2014). Targeting c-Met in melanoma: Mechanism of resistance and efficacy of novel combinatorial inhibitor therapy. Cancer Biol. Ther. 15, 1129–1141.

21. Evans, E.K., Tester, R., Aslanian, S., Karp, R., Sheets, M., Labenski, M.T., Witowski, S.R., Lounsbury, H., Chaturvedi, P., Mazdiyasni, H., et al. (2013). Inhibition of Btk with CC-292 provides early pharmacodynamic assessment of activity in mice and humans. J. Pharmacol. Exp. Ther. 346, 219–228.

22. Fan, L., Young, P.R., Barone, F.C., Feuerstein, G.Z., Smith, D.H., and McIntosh, T.K. (1996). Experimental brain injury induces differential expression of tumor necrosis factor-α mRNA in the CNS. Mol. Brain Res. 36, 287–291.

23. Fourgeaud, L., Traves, P.G., Tufail, Y., Leal-Bailey, H., Lew, E.D., Burrola, P.G., Callaway, P., Zagorska, A., Rothlin, C. V., Nimmerjahn, A., et al. (2016). TAM receptors regulate multiple features of microglial physiology. Nature 532, 240–244.

24. Frugier, T., Morganti-Kossmann, M.C., O’Reilly, D., and McLean, C.A. (2010). In situ detection of inflammatory mediators in post mortem human brain tissue after traumatic injury. J. Neurotrauma 27, 497–507.

25. Gao, C.F., and Woude, G.F.V. (2015). The MET receptor family. In Receptor Tyrosine Kinases: Family and Subfamilies, (Springer International Publishing), pp. 321–358.

26. Gordin, M., Tesio, M., Cohen, S., Gore, Y., Lantner, F., Leng, L., Bucala, R., and Shachar, I. (2010). c-Met and Its Ligand Hepatocyte Growth Factor/Scatter Factor Regulate Mature B Cell Survival in a Pathway Induced by CD74. J. Immunol. 185, 2020–2031.

27. Halleskog, C., Dijksterhuis, J.P., Kilander, M.B.C., Becerril-Ortega, J., Villaescusa, J.C., Lindgren, E., Arenas, E., and Schulte, G. (2012). Heterotrimeric G protein-dependent WNT-5A signaling to ERK1/2 mediates distinct aspects of microglia proinflammatory transformation. J. Neuroinflammation 9.

28. Herzog, C., Garcia, L.P., Keatinge, M., Greenald, D., Moritz, C., Peri, F., and Herrgen, L. (2019). Rapid clearance of cellular debris by microglia limits secondary neuronal cell death after brain injury in vivo. Dev. 146.

29. Holschneider, D.P., Guo, Y., Wang, Z., Vidal, M., and Scremin, O.U. (2019). Positive allosteric modulation of cholinergic receptors improves spatial learning after cortical contusion injury in Mice. J. Neurotrauma 36, 2233–2245.

30. Hopp, S., Nolte, M.W., Stetter, C., Kleinschnitz, C., Sirén, A.L., and Albert-Weissenberger, C. (2017). Alleviation of secondary brain injury, posttraumatic inflammation, and brain edema formation by inhibition of factor XIIa. J. Neuroinflammation 14.

31. Hosonuma, M., Sakai, N., Furuya, H., Kurotaki, Y., Sato, Y., Handa, K., Dodo, Y., Ishikawa, K., Tsubokura, Y., Negishi-Koga, T., et al. (2021). Inhibition of hepatocyte growth factor/c-Met signalling abrogates joint destruction by suppressing monocyte migration in rheumatoid arthritis. Rheumatol. (United Kingdom) 60, 408–419.

32. Hu, X., Hu, X., Li, S., Li, S., Doycheva, D.M., Huang, L., Huang, L., Lenahan, C., Lenahan, C., Liu, R., et al. (2020). Rh-CSF1 attenuates neuroinflammation via the CSF1R/PLCG2/PKCL pathway in a rat model of neonatal HIE. J. Neuroinflammation 17.

33. Huang, L., Jiang, S., and Shi, Y. (2020). Tyrosine kinase inhibitors for solid tumors in the past 20 years (2001–2020). J. Hematol. Oncol. 13, 143.

34. Ito, M., Shichita, T., Okada, M., Komine, R., Noguchi, Y., Yoshimura, A., and Morita, R. (2015). Bruton’s tyrosine kinase is essential for NLRP3 inflammasome activation and contributes to ischaemic brain injury. Nat. Commun. 6.

35. Jassam, Y.N., Izzy, S., Whalen, M., McGavern, D.B., and El Khoury, J. (2017). Neuroimmunology of Traumatic Brain Injury: Time for a Paradigm Shift. Neuron 95, 1246–1265.

36. Kong, L.J., Li, H., Du, Y.J., Pei, F.H., Hu, Y., Zhao, L.L., and Chen, J. (2017). Vatalanib, a tyrosine kinase inhibitor, decreases hepatic fbrosis and sinusoidal capillarization in CCl4-induced fbrotic mice. Mol. Med. Rep. 15, 2604–2610.

37. Kyyriäinen, J., Ekolle Ndode-Ekane, X., and Pitkänen, A. (2017). Dynamics of PDGFRβ expression in different cell types after brain injury. Glia 65, 322–341.

38. Lai, W., Ouyang, L., Liu, N., Liu, S., Shi, Y., Chen, L., Wang, X., Xiao, D., Liu, S., and Tao, Y. (2020). Identification of CpG methylation and non-CpG methylation is associate with Lymphocyte specific helicase.

39. Lolkema, M.P., Bohets, H.H., Arkenau, H.T., Lampo, A., Barale, E., De Jonge, M.J.A., Van Doorn, L., Hellemans, P., De Bono, J.S., and Eskens, F.A.L.M. (2015). The c-Met tyrosine kinase inhibitor JNJ-38877605 causes renal toxicity through species-specific insoluble metabolite formation. Clin. Cancer Res. 21, 2297–2304.

40. Ma, M.W., Wang, J., Dhandapani, K.M., Wang, R., and Brann, D.W. (2018). NADPH oxidases in traumatic brain injury – Promising therapeutic targets? Redox Biol. 16, 285– 293.

41. Mathis, M. W., Mathis, A., & Uchida, N. (2017). Somatosensory cortex plays an essential role in forelimb motor adaptation in mice. Neuron, 93(6), 1493–1503.

42. Mecha, M., Yanguas-Casás, N., Feliú, A., Mestre, L., Carrillo-Salinas, F.J., Riecken, K., Gomez-Nicola, D., and Guaza, C. (2020). Involvement of Wnt7a in the role of M2c microglia in neural stem cell oligodendrogenesis. J. Neuroinflammation 17.

43. Moransard, M., Sawitzky, M., Fontana, A., and Suter, T. (2010). Expression of the HGF receptor c-met by macrophages in experimental autoimmune encephalomyelitis. Glia 58, 559–571.

44. Muyllaert, D., Terwel, D., Kremer, A., Sennvik, K., Borghgraef, P., Devijver, H., Dewachter, I., and Van Leuven, F. (2008). Neurodegeneration and neuroinflammation in cdk5/p25-inducible mice: A model for hippocampal sclerosis and neocortical degeneration. Am. J. Pathol. 172, 470–485.

45. Nam, H.Y., Nam, J.H., Yoon, G., Lee, J.Y., Nam, Y., Kang, H.J., Cho, H.J., Kim, J., and Hoe, H.S. (2018). Ibrutinib suppresses LPS-induced neuroinflammatory responses in BV2 microglial cells and wild-type mice. J. Neuroinflammation 15, 271.

46. Nimmerjahn, A., Kirchhoff, F., and Helmchen, F. (2005). Neuroscience: Resting microglial cells are highly dynamic surveillants of brain parenchyma in vivo. Science (80-.). 308, 1314–1318.

47. Nishikoba, N., Kumagai, K., Kanmura, S., Nakamura, Y., Ono, M., Eguchi, H., Kamibayashiyama, T., Oda, K., Mawatari, S., Tanoue, S., et al. (2020). HGF-MET Signaling Shifts M1 Macrophages Toward an M2-Like Phenotype Through PI3K-Mediated Induction of Arginase-1 Expression. Front. Immunol. 11.

48. O’Donnell, S.L., Frederick, T.J., Krady, J.K., Vannucci, S.J., and Wood, T.L. (2002). IGF-I and microglia/macrophage proliferation in the ischemic mouse brain. Glia 39, 85–97.

49. O’Keeffe, E., Kelly, E., Liu, Y., Giordano, C., Wallace, E., Hynes, M., Tiernan, S., Meagher, A., Greene, C., Hughes, S., et al. (2020). Dynamic Blood-Brain Barrier Regulation in Mild Traumatic Brain Injury. J. Neurotrauma 37, 347–356.

50. Organ, S.L., and Tsao, M.S. (2011). An overview of the c-MET signaling pathway. Ther. Adv. Med. Oncol. 3, S7–S19.

51. Pavlides, C., Miyashita, E. I. Z. O., & Asanuma, H. (1993). Projection from the sensory to the motor cortex is important in learning motor skills in the monkey. Journal of neurophysiology, 70(2), 733–741.

52. Pellerin, K., Rubino, S. J., Burns, J. C., Smith, B. A., McCarl, C. A., Zhu, J., et al. (2021). MOG autoantibodies trigger a tightly-controlled FcR and BTK-driven microglia proliferative response. Brain.

53. Ray, A.K., DuBois, J.C., Gruber, R.C., Guzik, H.M., Gulinello, M.E., Perumal, G., Raine, C., Kozakiewicz, L., Williamson, J., and Shafit-Zagardo, B. (2017). Loss of Gas6 and Axl signaling results in extensive axonal damage, motor deficits, prolonged neuroinflammation, and less remyelination following cuprizone exposure. Glia 65, 2051–2069.

54. Reardon, D.A., Egorin, M.J., Desjardins, A., Vredenburgh, J.J., Beumer, J.H., Lagattuta, T.F., Gururangan, S., Herndon, J.E., Salvado, A.J., and Friedman, H.S. (2009). Phase I pharmacokinetic study of the vascular endothelial growth factor receptor tyrosine kinase inhibitor vatalanib (PTK787) plus imatinib and hydroxyurea for malignant glioma. Cancer 115, 2188–2198.

55. Di Renzo, M.F., Bertolotto, A., Olivero, M., Putzolu, P., Crepaldi, T., Schiffer, D., Pagni, C.A., and Comoglio, P.M. (1993). Selective expression of the Met/HGF receptor in human central nervous system microglia. Oncogene 8, 219–222.

56. Ritzel, R.M., He, J., Li, Y., Cao, T., Khan, N., Shim, B., Sabirzhanov, B., Aubrecht, T., Stoica, B.A., Faden, A.I., et al. (2021). Proton extrusion during oxidative burst in microglia exacerbates pathological acidosis following traumatic brain injury. Glia 69, 746–764.

57. Roselli, F., Chandrasekar, A., and Morganti-Kossmann, M.C. (2018). Interferons in traumatic brain and spinal cord injury: Current evidence for translational application. Front. Neurol. 9.

58. Roth, T.L., Nayak, D., Atanasijevic, T., Koretsky, A.P., Latour, L.L., and McGavern, D.B. (2014). Transcranial amelioration of inflammation and cell death after brain injury. Nature 505, 223–228.

59. Row, P.E., Clague, M.J., and Urbé, S. (2005). Growth factors induce differential phosphorylation profiles of the Hrs-STAM complex: A common node in signalling networks with signal-specific properties. Biochem. J. 389, 629–636.

60. Saijo, K., and Glass, C.K. (2011). Microglial cell origin and phenotypes in health and disease. Nat. Rev. Immunol. 11, 775–787.

61. Shah, V.B., Ozment-Skelton, T.R., Williams, D.L., and Keshvara, L. (2009). Vav1 and PI3K are required for phagocytosis of β-glucan and subsequent superoxide generation by microglia. Mol. Immunol. 46, 1845–1853.

62. Shankar, A., Borin, T.F., Iskander, A., Varma, N.R.S., Achyut, B.R., Jain, M., Mikkelsen, T., Guo, A.M., Chwang, W.B., Ewing, J.R., et al. (2016). Combination of vatalanib and a 20-HETE synthesis inhibitor results in decreased tumor growth in an animal model of human glioma. Onco. Targets. Ther. 9, 1205–1219.

63. Sierksma, A., Lu, A., Mancuso, R., Fattorelli, N., Thrupp, N., Salta, E., Zoco, J., Blum, D., Buée, L., De Strooper, B., et al. (2020). Novel Alzheimer risk genes determine the microglia response to amyloidLβ but not to TAU pathology. EMBO Mol. Med. 12.

64. Spiller, K.J., Restrepo, C.R., Khan, T., Dominique, M.A., Fang, T.C., Canter, R.G., Roberts, C.J., Miller, K.R., Ransohoff, R.M., Trojanowski, J.Q., et al. (2018). Microglia-mediated recovery from ALS-relevant motor neuron degeneration in a mouse model of TDP-43 proteinopathy. Nat. Neurosci. 21, 329–340.

65. Suh, H.-S., Kim, M.-O., and Lee, S.C. (2005). Inhibition of Granulocyte-Macrophage Colony-Stimulating Factor Signaling and Microglial Proliferation by Anti-CD45RO: Role of Hck Tyrosine Kinase and Phosphatidylinositol 3-Kinase/Akt. J. Immunol. 174, 2712–2719.

66. Suzuki, Y., Funakoshi, H., Machide, M., Matsumoto, K., and Nakamura, T. (2008). Regulation of cell migration and cytokine production by HGF-like protein (HLP)/macrophage stimulating protein (MSP) in primary microglia. Biomed. Res. 29, 77–84.

67. Szmydynger-Chodobska, J., Strazielle, N., Gandy, J.R., Keefe, T.H., Zink, B.J., Ghersi-Egea, J.F., and Chodobski, A. (2012). Posttraumatic invasion of monocytes across the blood-cerebrospinal fluid barrier. J. Cereb. Blood Flow Metab. 32, 93–104.

68. Terashima, T., Nakae, Y., Katagi, M., Okano, J., Suzuki, Y., and Kojima, H. (2018). Stem cell factor induces polarization of microglia to the neuroprotective phenotype in vitro. Heliyon 4, e00837.

69. Tondo, G., Perani, D., and Comi, C. (2019). TAM receptor pathways at the crossroads of neuroinflammation and neurodegeneration. Dis. Markers 2019, 1–13.

70. Tonge, P.J. (2018). Drug-Target Kinetics in Drug Discovery. ACS Chem. Neurosci. 9, 29–39.

71. Vidoni ED, Acerra NE, Dao E, Meehan SK, Boyd LA. Role of the primary somatosensory cortex in motor learning: An rTMS study. Neurobiol Learn Mem. 2010 May;93(4):532–9. doi: 10.1016/j.nlm.2010.01.011. Epub 2010 Feb 2. PMID: 20132902.

72. Wang, C.F., Zhao, C.C., Liu, W.L., Huang, X.J., Deng, Y.F., Jiang, J.Y., and Li, W.P. (2020). Depletion of microglia attenuates dendritic spine loss and neuronal apoptosis in the acute stage of moderate traumatic brain injury in mice. J. Neurotrauma 37, 43–54.

73. Wang, S., Gan, S., Yang, X., Li, T., Xiong, F., Jia, X., et al. (2021). Decoupling of structural and functional connectivity in hubs and cognitive impairment after mild traumatic brain injury. Brain Connectivity, (2021).

74. Welser-Alves, J. V., and Milner, R. (2013). Microglia are the major source of TNF-α and TGF-β1 in postnatal glial cultures; Regulation by cytokines, lipopolysaccharide, and vitronectin. Neurochem. Int. 63, 47–53.

75. Wicher, G., Wallenquist, U., Lei, Y., Enoksson, M., Li, X., Fuchs, B., Abu Hamdeh, S., Marklund, N., Hillered, L., Nilsson, G., et al. (2017). Interleukin-33 Promotes Recruitment of Microglia/Macrophages in Response to Traumatic Brain Injury. J. Neurotrauma 34, 3173–3182.

76. Willis, E.F., MacDonald, K.P.A., Nguyen, Q.H., Garrido, A.L., Gillespie, E.R., Harley, S.B.R., Bartlett, P.F., Schroder, W.A., Yates, A.G., Anthony, D.C., et al. (2020). Repopulating Microglia Promote Brain Repair in an IL-6-Dependent Manner. Cell 180, 833–846.e16.

77. Wood, J. M., Bold, G., Buchdunger, E., Cozens, R., Ferrari, S., Frei, J., et al. (2000). PTK787/ZK 222584, a novel and potent inhibitor of vascular endothelial growth factor receptor tyrosine kinases, impairs vascular endothelial growth factor-induced responses and tumor growth after oral administration. Cancer research, 60(8), 2178–2189.

78. Wu, S. Y., Pan, B. S., Tsai, S. F., Chiang, Y. T., Huang, B. M., Mo, F. E., & Kuo, Y. M. (2020). BDNF reverses aging-related microglial activation. Journal of Neuroinflammation, 17(1), 1–18.

79. Yamagata, T., Muroya, K., Mukasa, T., Igarashi, H., Momoi, M., Tsukahara, T., Arahata, K., Kumagai, H., and Momoi, T. (1995). Hepatocyte growth factor specifically expressed in microglia activated Ras in the neurons, similar to the action of neurotrophic factors. Biochem. Biophys. Res. Commun. 210, 231–237.

80. Yan EB, Satgunaseelan L, Paul E, Bye N, Nguyen P, Agyapomaa D, Kossmann T, Rosenfeld JV, Morganti-Kossmann MC. Post-traumatic hypoxia is associated with prolonged cerebral cytokine production, higher serum biomarker levels, and poor outcome in patients with severe traumatic brain injury. J Neurotrauma. 2014 Apr 1;31(7):618–29. doi: 10.1089/neu.2013.3087. Epub 2014 Jan 9. PMID: 24279428; PMCID: PMC3961772.

81. Yu, C.H., Nguyen, T.T.K., Irvine, K.M., Sweet, M.J., Frazer, I.H., and Blumenthal, A. (2014). Recombinant Wnt3a and Wnt5a elicit macrophage cytokine production and tolerization to microbial stimulation via Toll-like receptor 4. Eur. J. Immunol. 44, 1480–1490.

82. Zeisel, A., M□oz-Manchado, A.B., Codeluppi, S., Lönnerberg, P., Manno, G. La, Juréus, A., Marques, S., Munguba, H., He, L., Betsholtz, C., et al. (2015). Cell types in the mouse cortex and hippocampus revealed by single-cell RNA-seq. Science (80-.). 347, 1138–1142.

83. Zhang, J., Malik, A., Choi, H.B., Ko, R.W.Y., Dissing-Olesen, L., and MacVicar, B.A. (2014). Microglial CR3 activation triggers long-term synaptic depression in the hippocampus via NADPH oxidase. Neuron 82, 195–207.

84. Zhao, L., Wu, Y., Xie, X.D., Chu, Y.F., Li, J.Q., and Zheng, L. (2015). C-Met identifies a population of matrix metalloproteinase 9-producing monocytes in peritumoural stroma of hepatocellular carcinoma. J. Pathol. 237, 319–329.

85. Zhou, Z., Gao, S., Li, Y., Peng, R., Zheng, Z., Wei, W., Zhao, Z., Liu, X., Li, L., and Zhang, J. (2020). VEGI Improves Outcomes in the Early Phase of Experimental Traumatic Brain Injury. Neuroscience 438, 60–69.

