## Supplementary figures and images for "Met/HGFR triggers detrimental reactive microglia in TBI"

### supplementary figure 1

Figure_S1.pdf
2439 x 2705

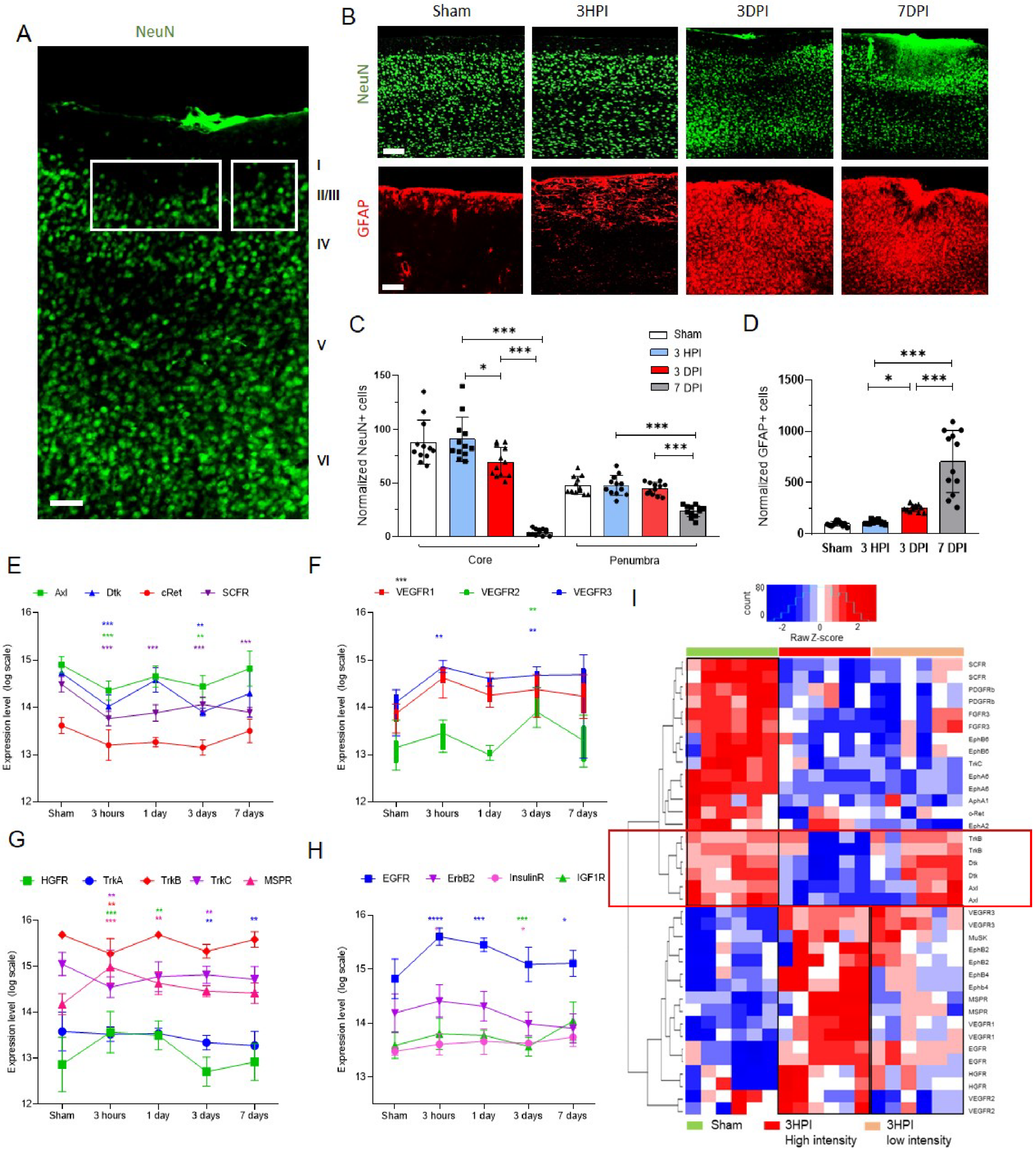

### supplementary figure 2

Figure_S2.pdf
2422 x 1910

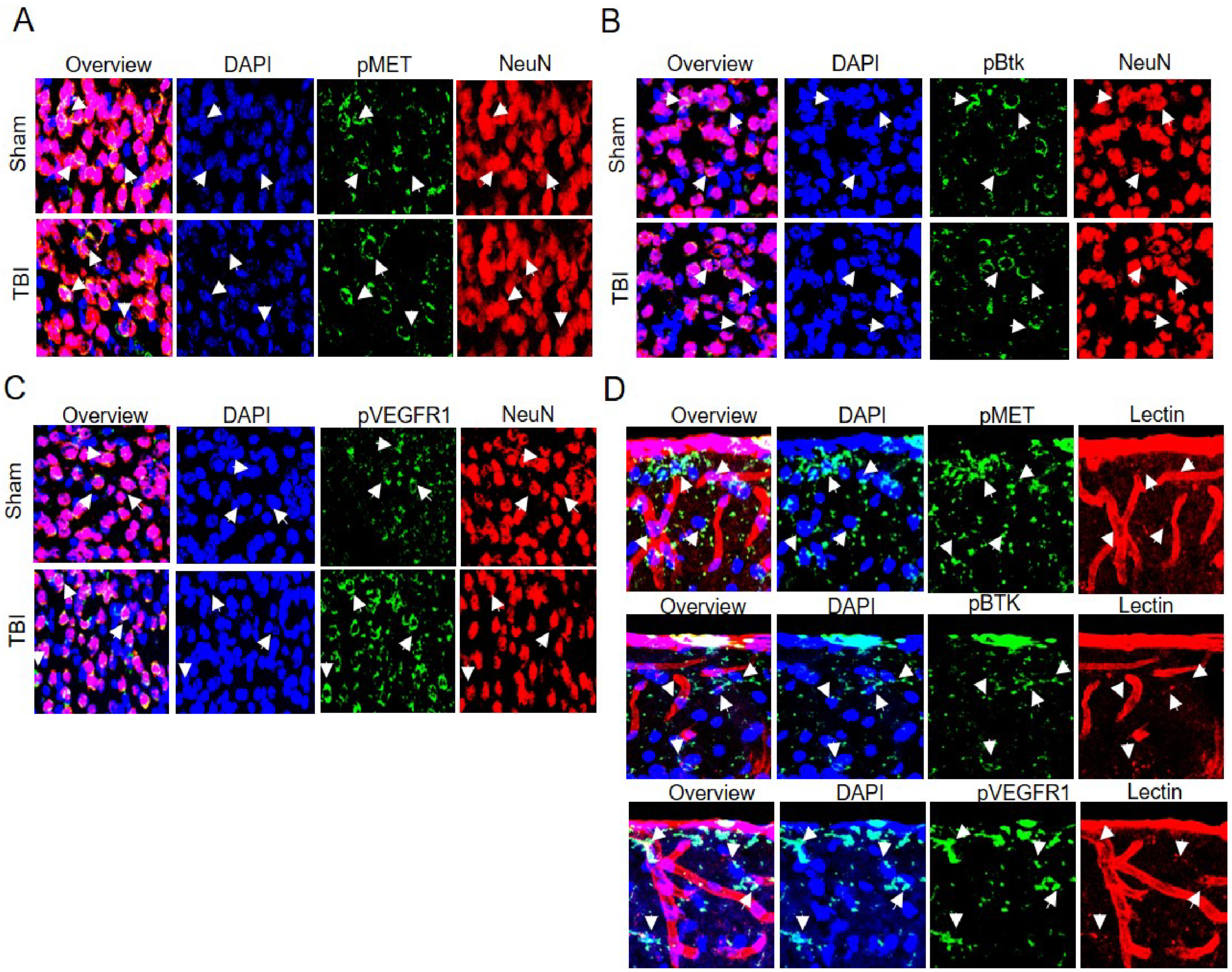

### supplementary figure 3

Figure_S3.pdf
1797 x 3405

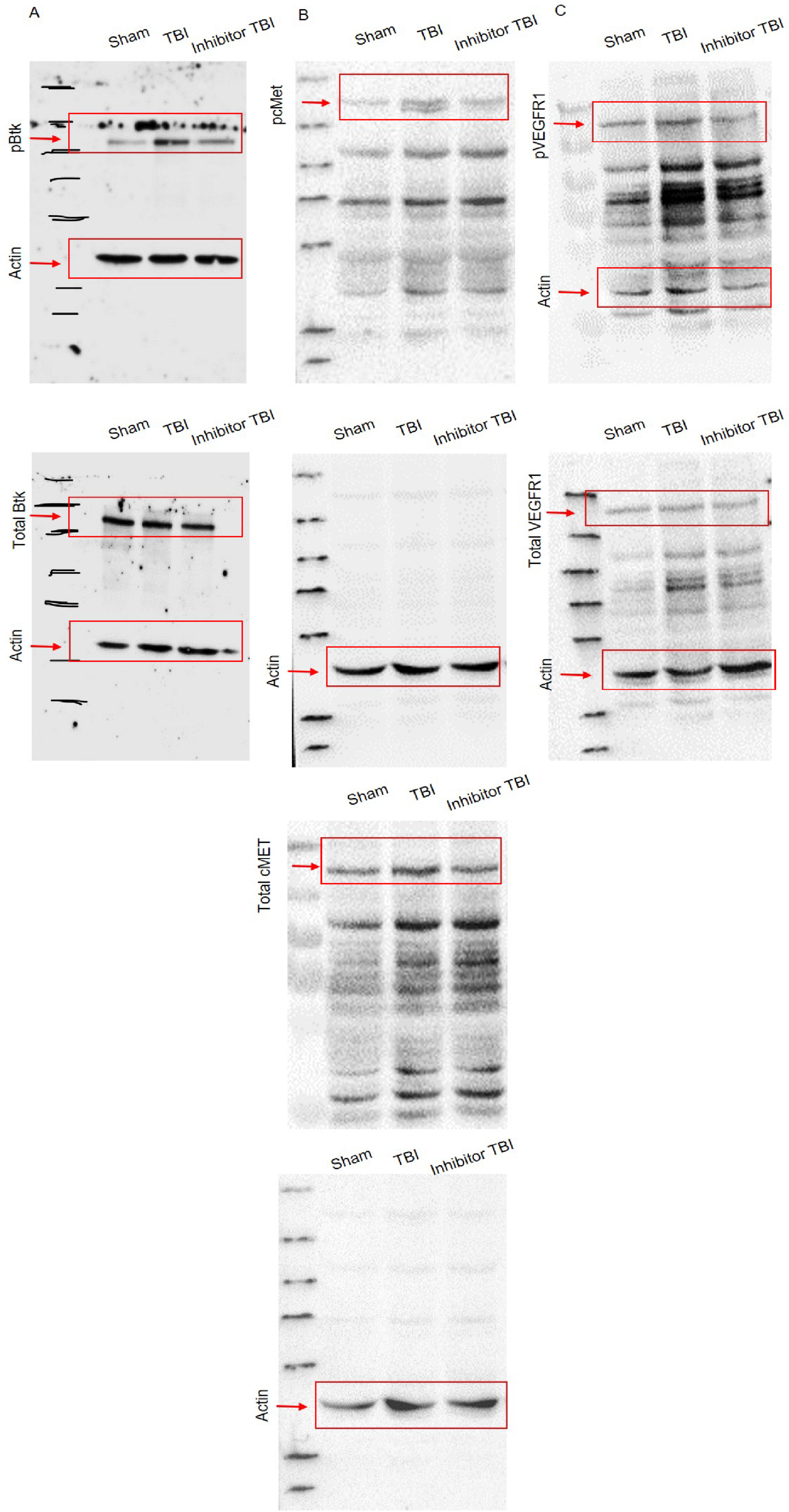

### supplementary figure 4

Figure_S4.pdf
2430 x 3217

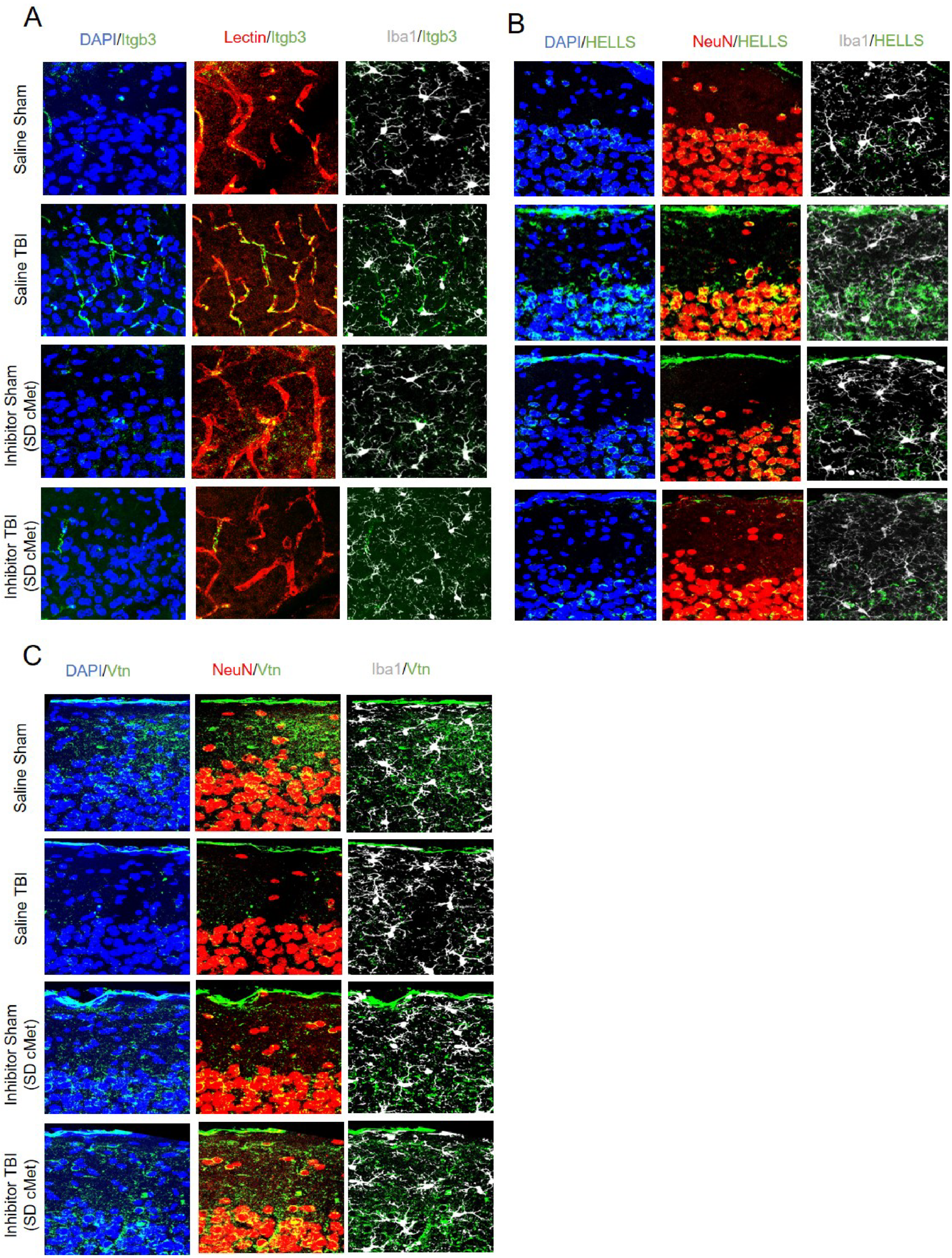
